# Dephosphorylation of 4EBP1/2 Induces Prenatal Neural Stem Cell Quiescence

**DOI:** 10.1101/2023.02.14.528513

**Authors:** Laura C. Geben, Asa A. Brockman, Mary Bronwen L. Chalkley, Serena R. Sweet, Julia E. Gallagher, Alexandra L. Scheuing, Richard B. Simerly, Kevin C. Ess, Jonathan M. Irish, Rebecca A. Ihrie

**Author notes:** **Contributions** LCG: conceptualization, data acquisition, funding acquisition, data analysis & interpretation, writing — original drafts & final version AAB: data analysis MBLC: data acquisition SRS: data acquisition, data analysis JEG: data analysis ALS: data acquisition RBS: supervision KCE: conceptualization, supervision, funding acquisition JMI: supervision, data analysis RAI: conceptualization, supervision, funding acquisition, data analysis & interpretation, writing - original drafts & final version All authors: review & approval of manuscript.

## Abstract

A limiting factor in the regenerative capacity of the adult brain is the abundance and proliferative ability of neural stem cells (NSCs). Adult NSCs are derived from a subpopulation of embryonic NSCs that temporarily enter quiescence during mid-gestation and remain quiescent until postnatal reactivation. Here we present evidence that the mechanistic/mammalian target of rapamycin (mTOR) pathway regulates quiescence entry in embryonic NSCs of the developing forebrain. Throughout embryogenesis, two downstream effectors of mTOR, p-4EBP1/2 T37/46 and p-S6 S240/244, were mutually exclusive in NSCs, rarely occurring in the same cell. While 4EBP1/2 was phosphorylated in stem cells undergoing mitosis at the ventricular surface, S6 was phosphorylated in more differentiated cells migrating away from the ventricle. Phosphorylation of 4EBP1/2, but not S6, was responsive to quiescence induction in cultured embryonic NSCs. Further, inhibition of p-4EBP1/2, but not p-S6, was sufficient to induce quiescence. Collectively, this work offers new insight into the regulation of quiescence entry in embryonic NSCs and, thereby, correct patterning of the adult brain. These data suggest unique biological functions of specific posttranslational modifications and indicate that the preferential inhibition of such modifications may be a useful therapeutic approach in neurodevelopmental diseases where NSC numbers, proliferation, and differentiation are altered.

## Introduction

The ventricular-subventricular zone (V-SVZ) is the largest neural stem cell (NSC) niche in the postnatal mammalian brain and cells from this niche produce multiple subtypes of neurons and glia (Alvarez-Buylla & García-Verdugo, 2002; Bond et al., 2021; Chaker et al., 2016; David-Bercholz et al., 2021; Delgado et al., 2021; Doetsch et al., 1997; Ihrie & Álvarez-Buylla, 2011; Merkle et al., 2004; Obernier & Alvarez-Buylla, 2019; Radecki & Samanta, 2022; Young et al., 2007). Postnatal adult NSCs, also termed B1 cells, are derived from a subpopulation of embryonic NSCs, termed pre-B1 cells, that enter a transient quiescence during mid-to late-neurogenesis, with the cell cycle slowing down beginning at embryonic day 13.5 and being completed by embryonic day 15.5. These cells remain quiescent until reactivation in early adulthood (postnatal days 21 — 28) (Fuentealba et al., 2015; Furutachi et al., 2015). Disruptions to either the cycling kinetics or total number of quiescent NSC populations in the embryo alter postnatal neurogenesis in both the V-SVZ and the dentate gyrus, the second major neurogenic niche (Berg et al., 2019; Bond et al., 2021; Hu et al., 2017; Kokovay et al., 2012; D. Y. Wang et al., 2020). While prior work has established that the regenerative capacity of the adult brain is directly related to pre-B1 cells’ transient quiescence entry, little is known about the dynamics and regulation of this essential process.

The mechanistic/mammalian target of rapamycin (mTOR) kinase is a principal regulator of cell growth (Laplante & Sabatini, 2012; G. Y. Liu & Sabatini, 2020) that has been shown to regulate the quiescence of other stem cell populations, including in postnatal NSCs (Cho & Hwang, 2012; Nieto­Gonzalez et al., 2019; Rodgers et al., 2014; Rossi et al., 2021; Sousa-Nunes et al., 2011). However, this potential relationship has not yet been explored in embryonic NSCs. The two primary downstream effectors of mTOR, phosphorylated ribosomal S6 kinase (p-S6) and phosphorylated 4E-binding proteins (p-4EBP1/2), have not been systematically characterized together in embryonic NSCs throughout neurogenesis or in the context of quiescence. Canonically, p-S6 regulates cell size and growth through its function in the ribosome, while p-4EBP1/2 is involved in cap-dependent mRNA translation (Gingras et al., 1999; Magnuson et al., 2012; Montagne et al., 1999; Ruvinsky & Meyuhas, 2006). However, in many studies, either p-S6 is used as the sole readout of mTOR activity, or the two proteins are used interchangeably. A growing body of evidence suggests that these two proteins have distinct, non-compensatory biological functions that are triggered by phosphorylation (Magnuson et al., 2012). Specifically, in postnatal NSCs, p-4EBP1/2 has been shown to regulate selective translation and regulation of self-renewal (Hartman et al., 2013). In glioblastoma cell lines, only p-4EBP1/2 was shown to correlate with a cell’s entry to or exit from the cell cycle (Fan et al., 2017, 2018).

Additionally, p-4EBP1/2 and p-S6 can behave independently in response to different ligands binding the upstream receptors that can activate this signaling pathway, such as insulin, growth factors, or amino acids (Sparta et al., 2021). Establishing the differences in activity and function between p-S6 and p-4EBP1/2 in embryonic NSCs may have direct relevance for patients with disorders of dysregulated mTOR signaling for whom mTOR inhibitors are often prescribed (Cavalheiro et al., 2021; Ebrahimi-Fakhari et al., 2021; Franz, 2011; Karalis & Bateup, 2021; Overwater et al., 2019). Three generations of mTOR inhibitors currently exist and have differing potencies and relative abilities to inhibit p-S6 and p­4EBP1/2. First generation rapamycin and analog drugs, termed “rapalogs,” have repeatedly been shown to have a greater effect on p-S6 than p-4EBP1/2 and on mTOR complex 1 (mTORC1) than complex 2 (mTORC2) (Choi et al., 1996; Fan et al., 2018). Second generation Tork inhibitors were designed to have improved potency against mTORC2 (Feldman et al., 2009), while third generation inhibitors, including RapaLink-1, were designed to have improved potency against p-4EBP1/2 (Rodrik-Outmezguine et al., 2016). This array of inhibitors offers an opportunity to separate the biological effects of p-S6 and p­4EBP1/2 in embryonic NSCs, where they have been untested, and their potential contributions to quiescence.

Here, the signaling patterns of mTOR targets in embryonic NSCs of the V-SVZ and their potential involvement in pre-B1 cell quiescence were investigated by adapting an established *in vitro* model of reversible quiescence to prenatal cells and quantifying functional effects of modulating downstream targets of mTORC1. p-4EBP1/2 and p-S6 were found to be expressed independently, not coordinately, in distinct populations of NSCs in the embryonic brain. The proliferative ability of an embryonic NSC was dependent upon phosphorylation of 4EBP1/2, as decreases in p-4EBP1/2, but not S6, were sufficient to induce quiescence. These results suggest mTOR-dependent phosphorylation of 4EBP1/2 is a key regulatory step in quiescence entry of embryonic pre-B1 NSCs and thus establishment of the postnatal stem cell niche.

## Materials and Methods

### Animals

All procedures involving animals were performed in accordance with animal health, safety, and wellness protocols outlined by both institutional (Institutional Animal Care and Use Committee) and national (National Institute of Health) governing bodies. Wild type C57 Black 6 mice were obtained from Charles River Laboratories. To collect embryos for cells, timed pregnant embryonic day 15.5 (E15.5) dams were euthanized via Avertin overdose and dissected. The embryos were collected for culture generation as described in the following sections. To collect embryos for tissue sections, timed pregnant E13.5, E15.5, and E17.5 dams were euthanized via Avertin overdose and transcardially perfused with 0.9% saline followed by 4% paraformaldehyde solution diluted in 0.2M phosphate buffer. Dissected embryos were processed further for imaging as described in the following sections.

### iDISCO+ Tissue Clearing

Whole E13.5 embryos were processed, stained, and cleared according to the immunolabeling-enabled three-dimensional imaging of solvent-cleared organs (iDISCO+) protocol described in Renier et al., 2014. Briefly, following perfusion of the pregnant dam, the embryos were drop fixed in 4% paraformaldehyde overnight at 4°C, and stored in PBS containing 0.1% sodium azide until staining. The embryos were dehydrated in six methanol washes (20%, 40%, 60%, 80%, and 100% at room temperature and a 100% wash chilled to 4°C), stored overnight in 66% dichloromethane in methanol at 4°C, washed twice with methanol (100%) at room temperature, and bleached overnight with 5% hydrogen peroxide in methanol at 4°C. The samples were rehydrated through washes in methanol (80%, 60%, 40%, 20%), 1X PBS, and two washes in buffer containing PBS and 0.2% TritonX-100 — one at room temperature and one overnight at 4°C. All washes, except those that lasted overnight, were 1 hour. Samples were incubated for 12 hours at 37°C with a permeabilization solution containing 10% PBS/0.2% TritonX-100/20% DMSO/2.3% glycine and incubated for 12 hours at 37°C with blocking solution containing 10% PBS/0.2% TritonX-100/6% normal donkey serum/10% DMSO. The samples were incubated for 24 hours with primary antibodies in buffer containing 10% PBS/0.2% Tween-20/0.1% 10 mg/mL Heparin/5% DMSO/3% normal donkey serum at 37°C, washed for 24 hours in buffer containing 10% PBS/0.2% Tween-20/0.1% 10 mg/mL Heparin at 37°C, incubated for 24 hours with Alexa Fluor secondary antibodies (Thermo Fisher Scientific) in buffer containing 10% PBS/0.2% Tween-20/0.1% 10 mg/mL Heparin/3% normal donkey serum at 37°C, and washed for an additional 24 hours in buffer containing 10% PBS/0.2% Tween-20/0.1% 10 mg/mL Heparin at 37°C. Antibodies are listed in Table 2. The samples were dehydrated through six hour-long washes in methanol (20%, 40%, 60%, 80%, 100%, 100%), incubated for 3 hours in 66% dichloromethane in methanol, washed twice in dichloromethane for 15 minutes, and cleared and stored in dibenzyl ether until imaging at 4X and 15X via SmartSPIM (LifeCanvas Technologies) light sheet fluorescence microscope. !marls Software version 9.5.1 was used for image and video reconstruction.

### Immunostaining of Mouse Brains

The heads were removed from E13.5, E15.5, and E17.5 embryos, drop fixed in 4% paraformaldehyde 2 hours (for 10 p.m thick sections) or overnight (for 50 p.m thick sections) at 4°C, and then sunk in 30% sucrose solution. Fixed heads were embedded into optimal cutting temperature compound (OCT) (Tissue-Tek, Sakura, 4583) before cryosectioning and mounting on Color Frost Plus microscope slides (Thermo Fisher Scientific, 12-550-16). Slides with OCT-embedded embryonic brain slices were removed from the freezer and allowed to acclimate to room temperature for 20 minutes in the chemical hood. Slides were washed three times in 1X PBS for 5 minutes, incubated in blocking solution containing PBS/1% normal donkey serum/1% BSA/0.1% Triton X-100 for 30 minutes at room temperature. Primary antibodies and primary-secondary antibody conjugates were applied to the slides overnight at 4°C. Antibodies are listed in Table 2. Slides were washed again three times in 1X PBS for 5 minutes and Alexa Fluor secondary antibodies (Thermo Fisher Scientific) were applied to the slides for approximately 1 hour at room temperature. Slides were washed one time with 1X PBS for 5 minutes. 4111,6-diamidino-2-phenylindole (DAPI) (diluted 1:10,000 in 1X PBS) was applied to the slides for 20 minutes at room temperature. The slides were washed 2 final times in 1X PBS for 5 minutes and rinsed with ddH20. Mowiol or Fluoromount-G (Electron Microscopy Sciences, 1798425) were used to mount coverslips (Fisher Scientific, 12-544-18P) and the slides were allowed to dry overnight. Slides were imaged on an LSM 710 Confocal Microscope (Zeiss) at specified magnifications and z-stacks at the Vanderbilt Cell Imaging Shared Resource and Zen Blue software (Zeiss) was used for image acquisition and reconstruction.

### Human Induced Pluripotent Stem Cell (hiPSC) Cell Culture

Two hiPSC lines (1) GM25256s from the Coriell Institute and (2) 77s from the Sahin lab (Sundberg et al., 2018) were cultured as previously described (Armstrong et al., 2017; Chalkley et al., 2022; Snow et al., 2020). In brief, iPSCs were grown as colonies on Matrigel (Corning, 354277) coated 6 well plates in mTeSR1 medium (StemCell Tech, 85850) at 37°C and 5% CO_2_. Culture media was replaced daily, and the cells were passaged with ReLSR (StemCell Tech, 05872) upon reaching confluency.

### Cortical Neurosphere Culture

iPSCs at 70% confluence were incubated with Accutase (Stem Cell Technologies, 07920) for 5 minutes at 37°C to dissociate cells. Cells were pelleted via centrifugation for 5 minutes at room temperature at 300 x *g.* 3 million cells were added to one well of an Aggrewell™800 (StemCell Tech, 34815) with neural induction media, containing 1:1 mixture DMEM/F12 GlutaMAX (Gibco, 10565-018) and Neurobasal (Gibco, 21103049), 0.5X N2 (Gibco, 17502048), 0.5X B27 with vitamin A (17504044), 2.5 μg/mL insulin (Life Technologies, 12585014), 0.75X Glutamax (Gibco, 35050061), 50 μM nonessential amino acids (Sigma, M7145), 50 μM 2-mercaptoethanol (Sigma, M6250), 50 U/mL penicillin-streptomycin (Gibco, 15140122), 10 μM SB431542 (Cayman Chemical, 13031) and 100 nM LDN-193189 (Tocris, 6053). Cells were allowed to aggregate overnight into a sphere while maintained at 37°C and 5% CO_2_. Neural induction media was replaced daily for 10 days. After 24 hours in culture, the neurospheres were transferred into well plates and maintained in suspension on an orbital shaker (95 rpm).

### Immunostaining of Neurospheres

All neurospheres were fixed by incubation in 4% paraformaldehyde for 15 minutes at 4°C. Fixed samples were blocked with blocking buffer containing PBS/1% normal donkey serum/1% BSA/0.1% Triton X-100 for 1 hour at room temperature. Primary antibodies were diluted in blocking buffer and then incubated overnight at 4°C. Alexa Fluor secondary antibodies (Thermo Fisher Scientific) were diluted in blocking buffer and then incubated 1 hour at room temperature in the dark. Antibodies are listed in Table 2. Hoechst (diluted 1:10,000 in 1X PBS) was applied to the slides for 20 minutes at room temperature. Images were acquired using a Prime 95B camera mounted on a Nikon spinning disk confocal microscope using a Plan Apo Lambda 20x objective lens at the Vanderbilt Nikon Center of Excellence. The software used for image acquisition and reconstruction were NIS-Elements Viewer (Nikon) and ImageJ (FIJI).

### Cell Pellet Preparation

Cultured cells were dissociated using Accutase and pelleted via centrifugation for 5 minutes at room temperature at 100 x *g* prior to fixation with 1.6% paraformaldehyde for 15 minutes, washed with 1x PBS, and re-pelleted via centrifugation for 5 minutes at room temperature at 100 x *g*. The supernatant was removed and replaced with 70% ethanol. The pellet was then paraffin embedded and prepared as 5-7 µm sections.

### Quantification of *ex vivo* Mouse Imaging Data

A custom Stardist 3D nuclear segmentation model was trained using 16 expert annotated cropped regions of interest from the dataset using the protocol described at https://github.com/stardist/stardist. Nuclear segmentation model was applied to each image in dataset followed by 3D pixel expansion to segment probable cell bodies. Marker mean intensity was measured and recorded for each nucleus, cytoplasm, and cell body (nucleus + cytoplasm). For each measured marker, cells were scored as positive or negative by thresholding. Thresholds were set and confirmed per staining batch by two independent viewers (LCG and AAB).

### Quantification of *in vitro* Human Imaging Data

A custom Stardist 2D nuclear segmentation model was trained using 18 expert annotated cropped regions of interest from the dataset using the protocol described at https://github.comistardist/stardist. The model was applied to each image in the dataset followed by 2D pixel expansion to segment probable cell bodies. Marker mean intensity was measured and recorded for each nucleus, cytoplasm, and cell body (nucleus + cytoplasm). For each measured marker, cells were scored as positive or negative by thresholding. Thresholds were set and confirmed per staining batch by two independent viewers (LCG and AAB).

### Primary Cell Cultures

Mouse embryos were collected at E15.5 from timed pregnant dams. To obtain stem cell cultures from the developing V-SVZ, a “top-down” dissection approach was employed. Briefly, the heads were separated from the body, the two brain hemispheres were pulled back to either side to reveal the developing cortex which was collected as dorsal NSCs. The developing ganglionic eminences were collected as ventral NSCs. The collected tissue was minced and incubated at 37°C, 5% CO2 with 0.25% trypsin-EDTA solution for 20 minutes while rocking. The tissue was then gently dissociated via trituration with a pipette and the cells were cultured and maintained in media specific to embryonic neural stem cells as described in Moghadam et al., 2018: Neurobasal media (ThermoFisher, 21103049); 1X B27 supplement without vitamin A (ThermoFisher, 12587010); 20 ng/mL mouse epidermal growth factor (ThermoFisher, 53003018); 10 ng/mL mouse basic fibroblast growth factor (ThermoFisher, PMG0035); 1 U/mL heparin (Sigma, 9041-08-01); 1X GlutaMax (ThermoFisher, 35050061); 1X modified Eagle’s medium non-essential amino acids (11140050); 0.1 mM [3-mercaptoethanol; 10 [ig/mL gentamicin. Cells were fed every 2-3 days and passaged upon reaching confluence.

### Quiescence Induction

To induce quiescence, embryonic NSC media was prepared as above without epidermal growth factor and with the addition of 50 ng/mL mouse bone morphogenetic protein 4 (R&D Systems, 5020-BP-010, stock dissolved in 4 mM hydrochloric acid with 0.1% BSA). Cells were fed every 2­3 days.

### Pharmacological modulators

mTOR inhibitors were added to culture media at concentrations listed in text and figure legends. Initial dissolutions of inhibitors were in DMSO, as specified by manufacturers’ instructions, and subsequent dilutions of concentrated stocks were in PBS.

**Table 1.**

| Reagent | Vendor | Catalog Number |
| --- | --- | --- |
| 4EGI-1 | Selleck Chem | S7369 |
| RapaLink-1 | Cell Signaling Technologies | 88626S |
| Rapamycin | Tocris | 1292 |
| Torin1 | Selleck Chem | S2827 |
| Torin2 | Selleck Chem | S2817 |
| Torkinib (PP242) | Selleck Chem | S2218 |

### Fluorescence flow cytometry

Cultured NSCs were collected for flow cytometry as previously described (Irish et al., 2010; Rushing et al., 2019). Briefly, cultures were gently dissociated for 7-10 minutes using Accutase at 37°C, pelleted by centrifuging for 5 minutes at room temperature at 100 x g, resuspended in 1 mL of the original spent media in 5 mL round bottom tubes (352052; Corning), and allowed to rest at 37°C for 1 hour and 25 minutes. Alexa Fluor 700-succinimidyl ester (Invitrogen, A20010) was added to the media at 37°C for 5 minutes to label non-intact cells. Samples that were to be live stained were then centrifuged for 5 minutes at room temperature at 100 x g, incubated with anti-VCAM1 antibody for 15 minutes at room temperature, rinsed with 1X PBS, and centrifuged again for 5 minutes at room temperature at 100 x g. All samples were then fixed with 1.6% paraformaldehyde, permeabilized with 70% ice cold ethanol, and stored at -20°C until staining. On the day of staining, samples were washed 2X with PBS and spun at 800 x g for 5 minutes at room temperature. Cells were stained with a cocktail of antibodies against intracellular antigens diluted in PBS/1% BSA for 30-60 minutes at room temperature. Samples were washed with PBS and spun at 800 x g for 5 minutes at room temperature. Samples were analyzed on either a Fortessa 4-laser or 5-laser instrument. Beads (Invitrogen, A10513) stained with a single fluorophore and unstained cells from the same cell line and treatment conditions were used for compensation and sizing controls. Gating was performed to isolate live, intact, single cells. Signaling values for each antigen were determined as both (1) the arcsinh transformed and (2) fold change of each sample compared to the column minimum for each antigen. The arcsinh scale used has been previously described (Irish et al., 2010; Rushing et al., 2019). On the arcsinh scale, a difference in 0.4 corresponds to a nearly 2-fold difference in total protein. All analyses were performed in Cytobank.

### Cell cycle analysis

Cultured cells were dissociated for 7-10 minutes using Accutase at 37°C, pelleted by centrifuging for 5 minutes at room temperature at 100 x g, resuspended in the original spent media in 5 mL round bottom tubes and incubated for 5 minutes with AlexaFluor700-succinimidyl ester to label non-intact cells. Cells were fixed with 1.6% paraformaldehyde, permeabilized with 70% ice cold ethanol, and stored at -20°C until staining. On the day of staining, samples were washed with PBS and spun at 800 x g for 5 minutes at room temperature twice. Cell pellets were resuspended in 1.5 [iN/1 Hoechst 33342 (Cell Signaling Technology, 4082) and incubated for 1 hour at 37°C and vortexed and every 15 minutes during incubation. Samples were analyzed on a BD LSRII 5-laser instrument. Cells in each phase of the cell cycle were determined via gating of live, intact, single cells. All analyses were performed in Cytobank.

### Quantification and statistical analysis

The quantification methods used and statistical tests performed are detailed in each figure and figure legend. GraphPad Prism 9 was used to perform all analyses.

### Data Availability Statement

All flow cytometry data will be made publicly available on FlowRepository upon publication (Spidlen et al., 2012).

### Antibodies

**Table 2.**

| **Antibody** | **Species** | **Clone** | **Catalog Number** | **Vendor** | **Dilution** |
| --- | --- | --- | --- | --- | --- |
| CD106 (VCAM1) conjugated to BV605 | Rat | 429 | 745193 | BD Biosciences | 1:200 |
| Ki67 | Rat | Sola15 | 14-5698-82 | Thermo Fisher | 1:200 |
| Ki67 conjugated to PerCP-Cy5.5 | Mouse | B56 | 561284 | BD Biosciences | 1:40 |
| p-4EBP1/2 T37/46 | Rabbit | 236B4 | 2855 | Cell Signaling Technologies | 1:800 (slides);<br>1:400 (iDISCO+) |
| p-4EBP1/2 T37/46 conjugated to Ax647 | Rabbit | 236B4 | 5123S | Cell Signaling Technologies | 1:100 (FC),<br>1:200 (IHC) |
| p-S6 S235/236 conjugated to Pacific Blue | Rabbit | D57.2.2E | 8520S | Cell Signaling Technologies | 1:100 |
| p-S6 S240/244 | Rabbit | D68F8 | 5364 | Cell Signaling Technologies | 1:800 (slides);<br>1:400 (iDISCO+) |
| p-S6 S240/244 conjugated to Ax488 | Rabbit | D68F8 | 5018S | Cell Signaling Technologies | 1:400 (Flow cytometry),<br>1:400 (IHC) |
| p-STAT3 S727 conjugated to PE | Mouse | 49P-STAT3 | 558557 | BD Biosciences | 1:15 |
| p-Vimentin S55 | Mouse | 4A4 | D076-3 | MBL | 1:1000 |
| Sox2 | Rat | Btce | 14-9811-82 | Invitrogen | 1:100 |
| Sox2 conjugated to PerCP-Cy5.5 | Mouse | O30-678 | 561506 | BD Biosciences | 1:400 |
| Tbr2 | Chicken | Polyclonal | AB15894 | Millipore | 1:250 |

## Experimental Design

### Mouse Experiments

The control groups for each experiment are listed in each figure legend. Both male and female embryos were used for immunohistochemistry experiments. Each embryo counted as a unique biological replicate (N=1). Embryos used in experiments presented here span at least two litters from each timepoint. Cultures were generated by pooling tissue collected from the indicated region from all the embryos (both male and female) from 1 or 2 timed pregnant dams, with an average litter size of 7. Cultures were grown up for one passage before cryopreservation. Each thawed cryovial counted as a unique biological replicate (N=1). Cultures used in experiments presented here represent three independent pregnant dam dissections and culture generations. Cultures treated with BMP4 were compared to matched cultures from the same cryovial that were not exposed to BMP4. Cultures treated with a pharmacological inhibitor were compared to matched cultures from the same cryovial treated with vehicle (1X PBS).

### Human Cells

Neurospheres were differentiated from two unique wild type induced pluripotent stem cell lines. Each differentiation counted as a unique biological replicate (N=1). Individual neurospheres from each biological replicate were considered technical replicates.

### Statistical Analysis

Unless otherwise indicated, all experiments had an N of at least 3. Exact Ns (separate biological replicates) are listed in the figure legends for each experiment. D’Agostino and Pearson K-squared tests were performed on each dataset to determine departure from normality. If a dataset failed to reach normality (p value < 0.05) or did not have enough replicates to perform the D’Agostino test (N < 8), then a nonparametric Mann-Whitney U test was performed. If the dataset reached normality (D’Agostino test p value > 0.05), a parametric student’s t-test was performed. Whether the test was paired or unpaired is indicated in the figure legends for each comparison. P values for each comparison are listed in the figure legends. GraphPad Prism 9 was used to perform all statistical analyses and generate plots.

## Results

### High levels of p-4EBP1/2, but not p-S6, are present in embryonic NSCs at the ventricle

To visualize levels of mTORC1 signaling prior to the genesis of pre-B1 cells, whole brains from embryonic day 13.5 mice were processed, cleared, and stained for p-S6 S240/244 and p-4EBP1/2 T37/46. p-S6 S240/244 was distributed in multiple cell layers of both the developing cortex and ganglionic eminences (Fig 1A). In contrast, p-4EBP1/2 T37/46 was primarily present in cell bodies lining the apical ventricular surface of developing cortex and was not abundant in cells that were more distant from the ventricle. In the lateral and medial ganglionic eminences, p-4EBP1/2 was similarly enriched at the ventricular surface and was also present in some cells deeper within the tissue (Fig 1B; full brain videos for p-S6 and p-4EBP1/2 in Extended Fig 1-1). To further investigate these patterns throughout neural development and the genesis of pre-B1 cells, corona! sections of embryonic day 13.5, 15.5, and 17.5 mouse brain were co-stained for both readouts of mTORC1 activity (Fig 1C). At all developmental ages measured, the two phosphorylated proteins were largely found in distinct cells. p-4EBP1/2 labeling was limited to cells lining the ventricular surface, many of which appeared to be actively undergoing mitosis (yellow arrows in Fig 1C). At E13.5 and 15.5, p-S6 was not found in cells lining the ventricular surface but was abundant in the subventricular zone and intermediate zones. At E17.5, p-S6 was observed in cells contacting the ventricular surface. The two phosphorylated proteins were rarely found in the same cell (E13: average 2.7% of counted cells [N=3], E15: average 0.52% of counted cells [N=4], E17: average 0.1% of counted cells [N=3]). The frequency of p-4EBP1/2-positive cells decreased with gestational age, from an average of 4.10% of cells per field at E13.5 to 0.17% at E17.5, while levels of p-S6 expression increased from 37.85% at E13.5 to 53.78% at E17.5.

**Figure 1:**
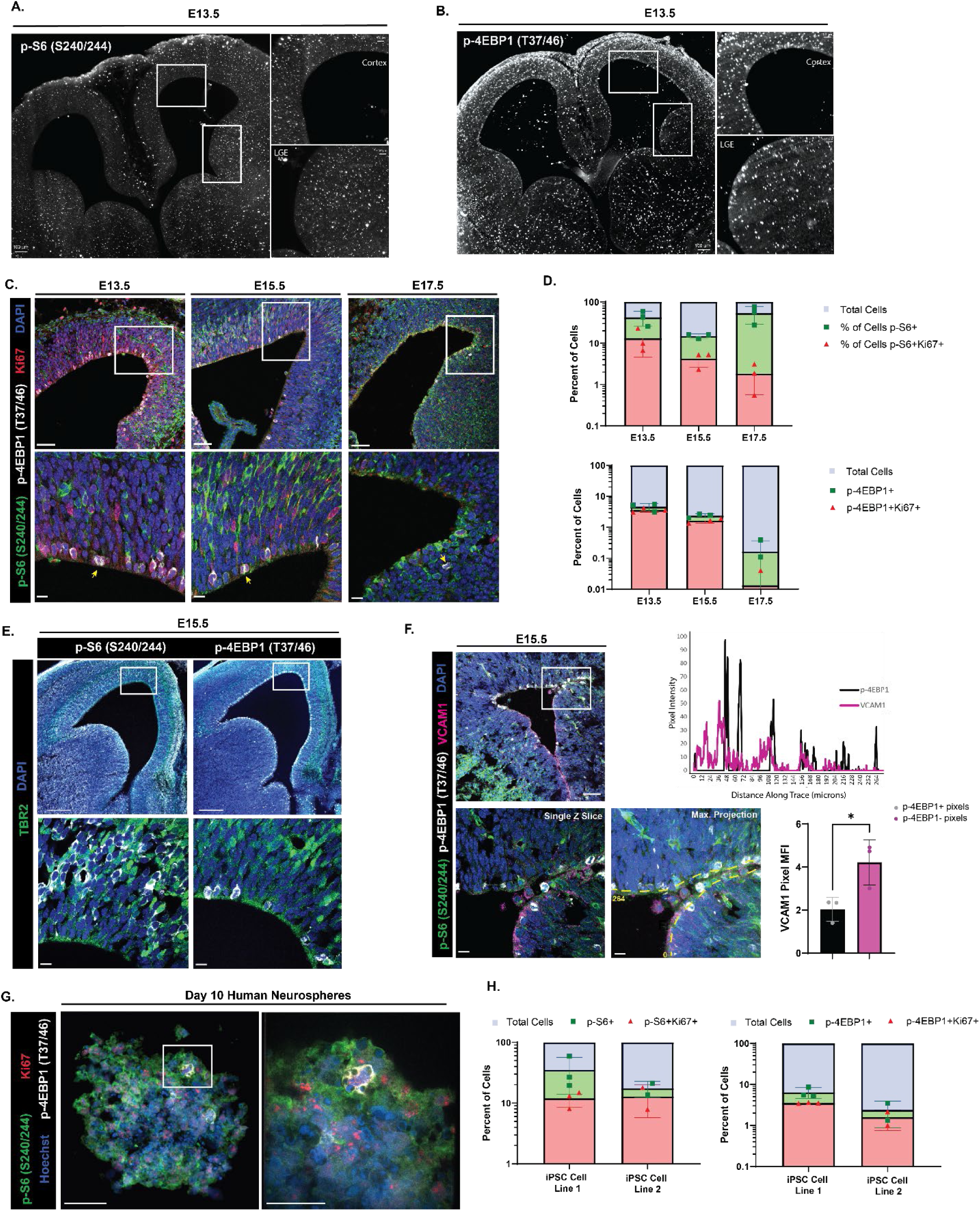
High levels of p-4EBP1/2, but not p-S6, are present in embryonic NSCs at the ventricle. (A) the serine 240/244 residues diffuse throughout the ventricular-subventricular zone tissue in both the developing cortex (inset, top) and lateral ganglionic eminence (inset, bottom). Scale bar information is listed on each figure. (B) iDISCO+-tissue cleared whole brain from an E13.5 mouse embryo (15X) showing phosphorylation of 4EBP1 at the threonine 37/46 residues limited to the cells immediately lining the ventricular surface and not present deeper into the subventricular zone in both the developing cortex (inset, top) and lateral ganglionic eminence (inset, bottom). Scale bar information is listed on each figure. (C) Staining of embryonic mouse brain for the downstream effectors of mTOR at E13.5, E15.5, and E17.5. 20X (top) representative images of the developing ventricular-subventricular zone with single slice of 63X z-stack (bottom) of the inset region showing p-S6 S240/244 (green), p-4EBP1 T37/46 (white), Ki67 (red), and DAPI (blue). Yellow arrows in the 63X representative images mark dividing cells with phosphorylated 4EBP1. Scale bars for all 20X images = 50 µm. Scale bars for all 63X images = 10 µm. (N for E13.5 = 3, N for E15.5 = 4, N for E17.5 = 3) (D) Quantification of the percent of shown in (C) across developmental time points. The percent of all cells positive for p-S6 S240/244 (green) and cells co-positive for p-S6 and Ki67 (red) (top). (2 way ANOVA with Tukey’s multiple comparisons test: percent of p-S6 positive cells: E13.5 versus E15.5 p = 0.0501, E15.5 versus E17.5 p = 0.0071, E13.5 versus E17.5 p = 0.5377; percent of p-S6 and Ki67 positive cells: E13.5 versus E15.5 p = 0.6704, E15.5 versus E17.5 p = 0.9697, E13.5 versus E17.5 p = 0.5301). The percent of all cells positive for p-4EBP1 T37/46 (green) and cells co-positive for p-4EBP1 and Ki67 (red) (bottom). (2 way ANOVA with Tukey’s multiple comparisons test: percent of p-4EBP1 positive cells: E13.5 versus E15.5 p = 0.0022, E15.5 versus E17.5 p = 0.0023, E13.5 versus E17.5 p < 0.0001; percent of p-4EBP1 and Ki67 positive cells: E13.5 versus E15.5 p = 0.0056, E15.5 versus E17.5 p = 0.0184, E13.5 versus E17.5 p < 0.0001). Error bars represent standard deviation. Plots showing statistics with significance shown in Extended Figure 1-1. (E) Staining of E15.5 mouse brain for the downstream effectors of mTOR and marker of intermediate progenitor cells. 5X representative images (top) of the developing ventricular-subventricular zone with single slice of 63X z-stack (bottom) of the inset region showing Tbr2 (green) colocalizing with p-S6 S240/244 (white, left) but not with p-4EBP1 T37/46 (white, right), and DAPI (blue). Scale bars for 5X images = 200 µm. Scale bars for 63X images = 10 µm. (F) Staining of E15.5 mouse brain for the downstream effectors of mTOR and protein required for radial glia maintenance and quiescence entry. 20X representative image (top left) of the developing ventricular-subventricular zone with single slice of 63X z-stack (bottom left) of the inset region showing p-S6 S240/244 (green), p-4EBP1 T37/46 (white), VCAM1 (magenta), and DAPI (blue). Line trace reporting pixel intensity for p-4EBP1 T37/46 (black) and VCAM1 (magenta) across distance of yellow dashed line shown in maximum projection image of the z stack for the inset region (top right). Maximum projection image of the z stack for the inset region (bottom middle). Quantification of the median fluorescence intensity of VCAM1 in pixels positive for p­4EBP1 (black) versus pixels negative for p-4EBP1 (magenta) (bottom right). Error bars represent standard deviation. (N = 3, paired two-tailed t test p value = 0.0175). Scale bars for 20X image = 50 µm. Scale bars for 63X images = 10 µm. (G) Staining of a wild type day 10 neurosphere derived from human induced pluripotent stem cells for the downstream effectors of mTOR. 40X (left) representative image of sphere within organoid with 100X image of the inset region (right) showing a dividing cell expressing p-S6 S240/244 (green), p-4EBP1 T37/46 (white), Ki67 (red), and Hoechst (blue). Scale bars for both images = 50 µm. (N = 2 representing 2 unique differentiations of 2 independent sets of wild type iPSC lines.) (H) Quantification of the percent of shown in (G) of day 10 cortical organoids. The percent of all cells positive for p-S6 S240/244 (green) and cells co-positive for p-S6 and Ki67 (red) (left). The percent of all cells positive for p-4EBP1 T37/46 (green) and cells co-positive for p-4EBP1 and Ki67 (red) (right). Error bars represent standard deviation.

Co-staining with Ki67, a marker of cycling cells, revealed that p-4EBP1/2 positive cells were often dividing: 80.97% of cells positive for p-4EBP1/2 at E13.5 were doubly positive for Ki67, 68.44% at E15.5, and 33.33% at E17.5; though Ki67 expression also decreased with age (27.02% of total cells at E13.5, 18.01% at E15.5, 2.54% at E17.5), consistent with prior reports (Hu et al., 2017) and the general decrease in neurogenesis at this stage (Fuentealba et al., 2015; Furutachi et al., 2015). p-S6 positive cells variably expressed Ki67 (31.26% of cells positive for p-S6 at E13.5, 27.93% at E15.5, 3.29% at E17.5). Additional Ki67 staining in the developing V-SVZ at E13.5 from cleared brain is shown in Extended Fig 1-1. To assign mTOR activity more precisely to radial glia and transit amplifying cells, tissues were co-stained for p-S6 S240/244, p-4EBP1/2 T37/46, and the transcription factor t-box brain protein 2 (Tbr2), which distinguishes the transit-amplifying progeny of cortical radial glia (Englund et al., 2005) (Fig 1E). Cells expressing Tbr2 were more likely to express p-S6 than p-4EBP1/2 at both E13.5 (12.99% Tbr2/p-S6 co-positive cells versus 0.65% Tbr2/p-4EBP1/2 co-positive cells) and E15.5 (34.72% Tbr2/p-S6 co-positive cells versus 2.26% Tbr2/p-4EBP1/2 co-positive cells).

As cells positive for p-4EBP1/2 were often co-positive for Ki67 and appeared to be actively dividing, this suggested that cells with low expression of p-4EBP1/2 could be pre-B1 cells, which have been reported to enter quiescence during prenatal development. Lineage tracing studies have indicated that pre-B1 cells begin entering quiescence as early as E13.5, with most completing quiescence entry by E15.5 (Fuentealba et al., 2015; Furutachi et al., 2015). E15.5 brains were co-stained with p-S6 S240/244, p-4EBP1/2 T37/46, and vascular cell adhesion molecule 1 (VCAM1), a protein required for the maintenance of the radial glia stem cell population and entry into quiescence (Hu et al., 2017; Kokovay et al., 2012; D. Y. Wang et al., 2020) (Fig 1F). VCAM1 was found infrequently in the developing cortex, but was more abundant in the ganglionic eminences, consistent with previously published findings (Hu et al., 2017) (Fig 1D, 20x). At E15.5, VCAM1 was expressed exclusively by cells at the ventricular surface and did not overlap with p-S6. Line scan analysis along the apical ventricular surface showed that cells positive for p-4EBP1/2 and VCAM1 were largely exclusive of each other (Figure 1D). In pixels negative for p-4EBP1/2, the average VCAM1 intensity was double that in pixels positive for p-4EBP1/2 (4.22 versus 2.04, N = 3).

To determine if similar patterns of p-4EBP1/2 / p-S6 dual abundance might be present in human neural progenitors, neurospheres were differentiated from two separate wild type human induced pluripotent stem cell lines (Sundberg et al., 2018). After 10 days in culture, neurospheres were collected and were co-stained with p-S6 5240/244, p-4EBP1/2 T37/46, and Ki67 (Fig 1G). As was observed in the mouse, p-S6 was more abundantly expressed than p-4EBP1/2 (21.78% of total cells positive for p-S6 versus 4.55% positive for p-4EBP1/2, N = 5 neurospheres) and cells positive for p-4EBP1/2 were often actively undergoing division and doubly positive for Ki67 (61.72% of p-4EBP1/2 positive cells). Similar to the pattern observed in the mouse, only a small percentage of cells (4.00% of total cells) were positive for both p-S6 and p-4EBP1/2.

### Exposure to BMP4 induces quiescence in embryonic NSCs *in vitro*

To determine how mTOR activity responds to changes in cellular proliferation and assess whether cells entering quiescence decrease levels of p-4EBP1/2, an *in vitro* model of quiescence was developed and validated. The role of bone morphogenetic protein 4 (BMP4) in directing differentiation and maintenance of stem cells in the V-SVZ and dentate gyrus has been extensively described (Li et al., 1998; Mira et al., 2010). BMP4 has been reported to induce a reversible quiescence *in vitro* in various stem cell populations within 72 hours, including postnatal neural stem cells (Knobloch et al., 2017; Mira et al., 2010; Rossi et al., 2021), but has not been tested in prenatal NSCs.

To test its use in prenatal NSC populations, neural stem cells dissected from the developing dorsal V-SVZ at embryonic day 15.5 were exposed to BMP4 with simultaneous withdrawal of EGF but maintenance of basic fibroblast growth factor (bFGF-2), and markers of cell proliferation and quiescence were quantified using fluorescence microscopy and flow cytometry. After 24 hours in media containing BMP4 and bFGF-2, levels of the proliferation marker Ki67 had significantly decreased compared to cells grown in control media containing bFGF-2 and epidermal growth factor (EGF) (Fig 2A), with a further decrease seen by 72 hours and persisting with additional time in BMP4 culture media (time course of quiescence entry shown in Extended Fig 2-1). Following extended incubation with BMP4/FGF-2 media for 6 days, Ki67 and p-vimentin levels both remained decreased compared to control (Fig 2B). BMP4-mediated decreases in Ki67 were reversible, as re-exposure to media containing EGF and lacking BMP4 rescued levels of Ki67 (Extended Fig 2-2). Within 24 hours of exposure to BMP4/FGF-2 media, levels of VCAM1 also began to rise compared to control cells and remained high at 72 hours (Fig 2C). To further verify that BMP4 was having the desired effect, the percentage of cells in each stage of the cell cycle was determined following 24 hours of exposure to BMP4. Cells grown in media containing BMP4/FGF-2 had an increased percentage of cells in the G0/G1 phase of the cell cycle compared to cells grown in media containing EGF/FGF-2 (averages of 80.83% vs 58.08%, N=9), and had a decreased percentage of cells in both the S (7.03% vs. 21.70%) and G2/M (8.98% vs 17.31%) phases of the cycle compared to control cells (Fig 2D; gates used to determine percent of cells in each phase of the cell cycle shown in Extended Fig 2-3).

**Figure 2:**
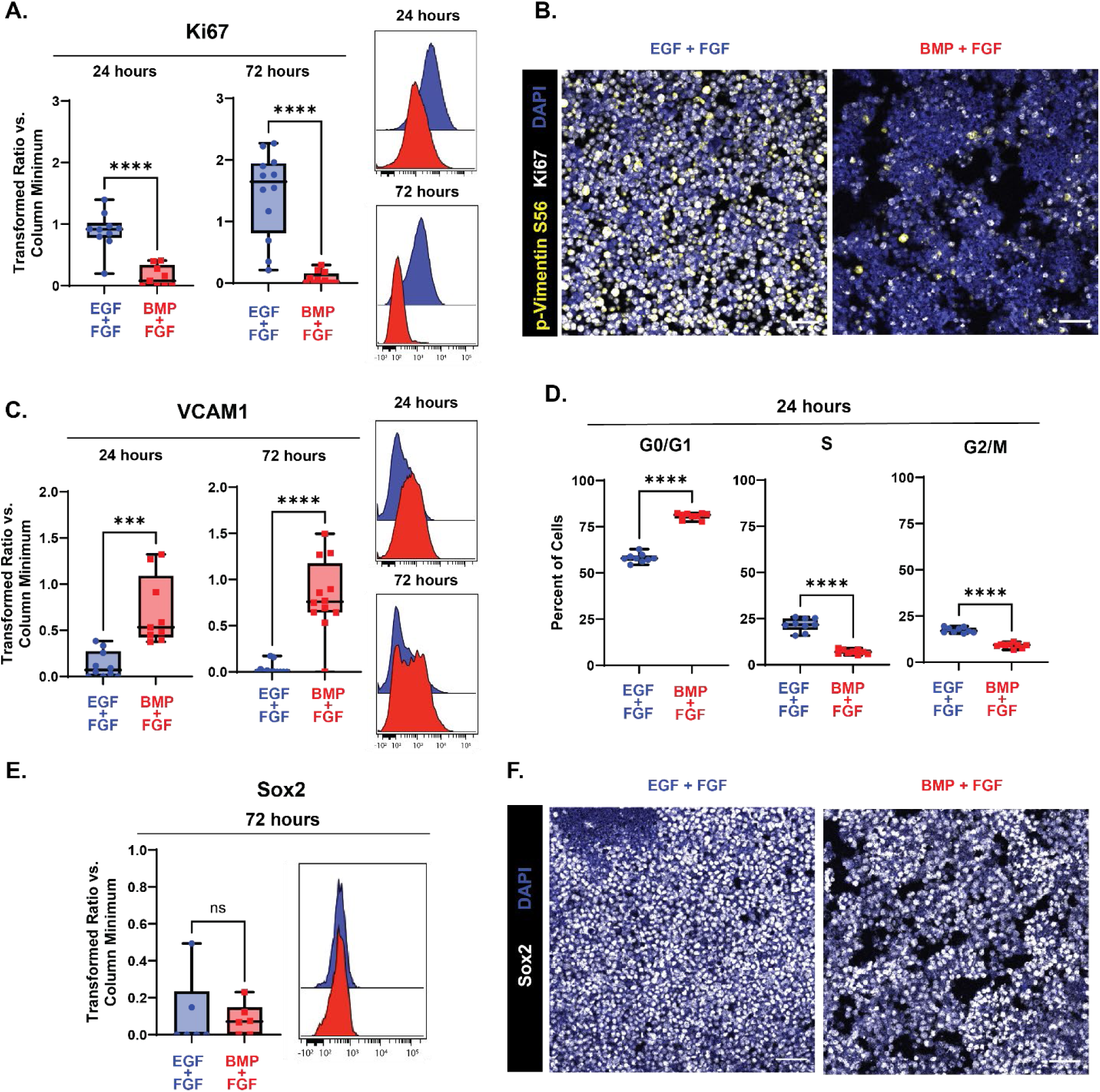
Exposure to BMP4 induces quiescence in embryonic NSCs *in vitro*. Quantification of proliferation and quiescence markers in E15.5 NSC cultures grown for 24 (left in plots) and 72 (right in plots) hours with media containing EGF/FGF (blue) or BMP4/FGF (red). For all plots, the Y axis depicts the arcsinh transformed ratio versus column minimum for a particular antigen. A difference of 0.4 corresponds to an approximately 2-fold difference of total protein. Representative histograms of 24 (top) and 72 hours (bottom). For all plots, error bars contact the maximum and minimum values. For Ki67 and VCAM1, at 24 hours N = 10 for EGF/FGF, N = 9 for BMP4/FGF; at 72 hours N = 12. (A) Quantification of levels of Ki67 (24 hours: unpaired two-tailed t-test p < 0.0001; 72 hours: unpaired two-tailed t-test p < 0.0001). (B) 20X representative images of E15.5 NSCs cultured in media with EGF/FGF (left) or BMP4/FGF (right) for p-vimentin S56 (yellow), Ki67 (white), and DAPI (blue). (C) Quantification of levels of VCAM1 (24 hours: unpaired two-tailed t-test p = 0.0003; 72 hours: unpaired two-tailed t-test p < 0.0001). (D) Percent of E15.5 NSC cultures grown for 24 hours with media containing EGF/FGF (blue) or BMP4/FGF (red) in each phase of the cell cycle. (N = 9, G0/G1 phase unpaired two-tailed t-test p < 0.0001, S phase unpaired two-tailed t-test p < 0.0001, G2/M unpaired two-tailed t-test p < 0.0001). (E) Quantification of levels of Sox2 (N = 6, Mann-Whitney test p = 0.6688). (F) 20X representative images of E15.5 NSCs cultured in media with EGF/FGF (left) or BMP4/FGF (right) stained for Sox2 (white) and DAPI (blue). Scale bars for all 20X images = 50 µm.

BMP4 has also been shown to direct NSCs into neuronal or astrocytic lineages depending on the time of expression (Katada et al., 2021); these effects that are directly antagonized and suppressed by EGF and FGF-2 to maintain stemness and self-renewal capacities (Lillien & Raphael, 2000; Sun et al., 2011). To ensure that the removal of EGF and exposure to BMP4 did not differentiation as reported for media containing only BMP4 without FGF-2, levels of Sox2, a transcription factor expressed by neural stem and progenitor cells, but not transit amplifying cells, were measured following acute (72 hours, Fig 2E) and prolonged (6 days, Fig 2F) exposure to BMP4. At 72 hours of BMP4 exposure, levels of Sox2 remained unchanged compared to cells grown in EGF/FGF-2 media. At 6 days in culture with BMP4, per-cell Sox2 expression remained high and comparable to cells grown in EGF/FGF-2 media. Further, cells cultured with BMP4 did not begin to express markers of differentiation, including TUJ1 or GFAP (Extended Fig 2-3). Taken together, these data indicate that consistent with published models, BMP4 in combination with FGF-2 efficiently induces reversible quiescence in embryonic NSCs *in vitro* within 24 hours without inducing differentiation.

### p-4EBP1/2 signaling decreases in embryonic NSCs following quiescence induction

To investigate whether low levels of p-4EBP1/2 were directly correlated with quiescence, neural stem cells dissected from the developing cortex at embryonic day 15.5 were cultured in media containing BMP4 and FGF-2 for acute (24 hours) or extended (72 hours) time periods and readouts of the mTOR pathway were measured. Two different phosphorylated sites on S6 were measured: S235/236, a residue phosphorylated by both the mTOR and MAPK/ERK pathways, and S240/244, a residue exclusively phosphorylated by the mTOR pathway (Magnuson et al., 2012; Roux et al., 2007). At both 24 and 72 hours in BMP4/FGF-2 media, neither of these readouts showed significant change relative to EGF/FGF-2 media (Figs 3B and Fig 3C), a pattern that was persisted after further prolonged exposure to BMP4/FGF-2 media (6 days, Fig 3D). In contrast, after 24 hours in media containing BMP4, levels of p-4EBP1/2 decreased compared to controls (Fig 3E), with a further decrease seen after 72 hours. To determine whether the decrease in p-4EBP1/2 remained after prolonged exposure to BMP4 and corresponded to a decrease in proliferation, cells were cultured for 6 days and co-stained for K167 and p-4EBP1/2 (Fig 3F). Levels of both Ki67 and p-4EBP1/2 decreased upon exposure to BMP4, compared to cells grown in EGF/FGF-2 media. At both 24 and 72 hours of BMP4 exposure, levels of an additional mTOR target, p-STAT3 S727 (Dodd et al., 2015; Yokogami et al., 2000), also decreased compared to the EGF/FGF-2 media condition (Fig 3A).

**Figure 3:**
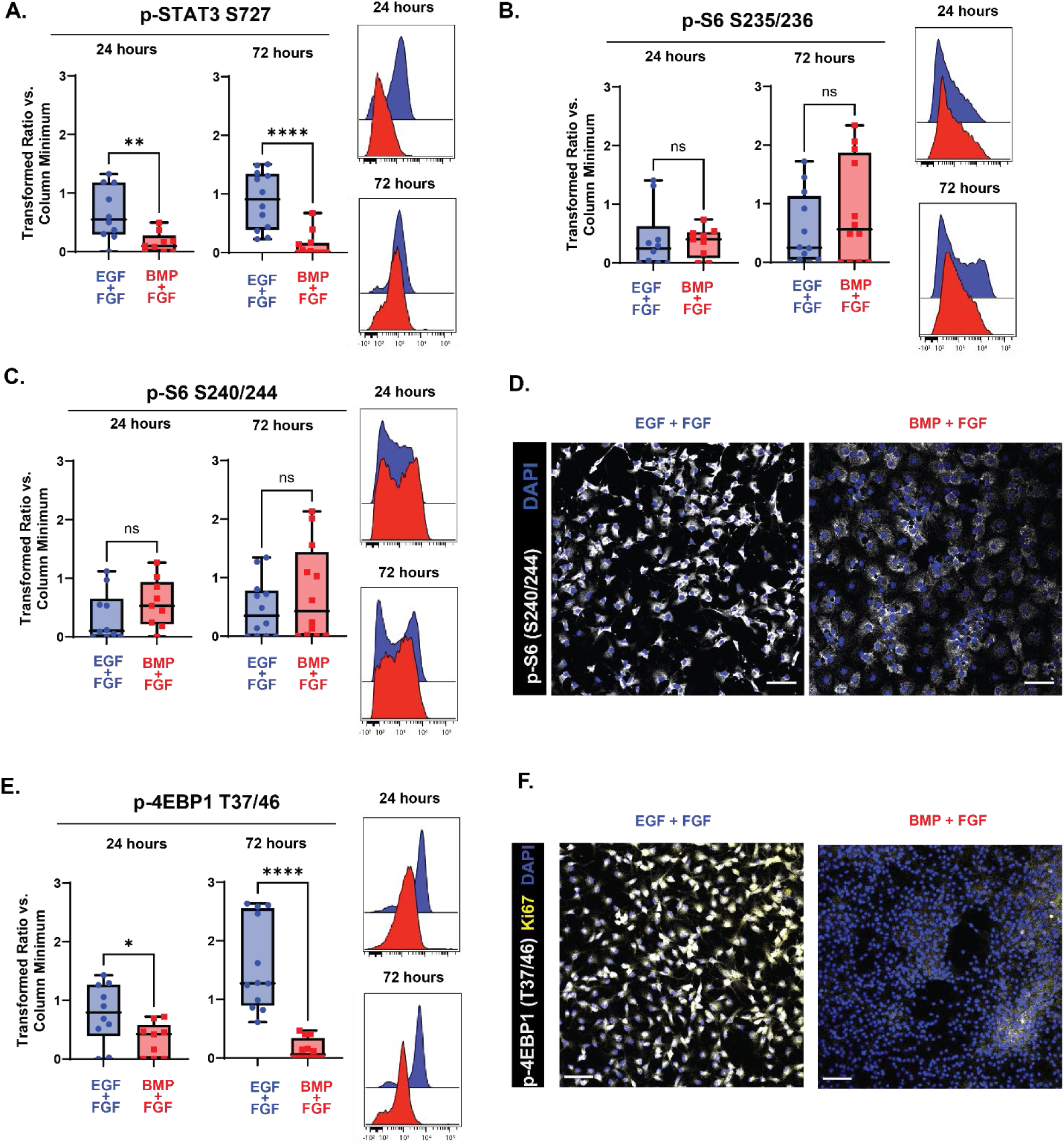
p-4EBP1/2 signaling decreases in embryonic NSCs following quiescence induction. Quantification of effectors downstream of mTOR in E15.5 NSC cultures grown for 24 (left in plots) and 72 (right in plots) hours with media containing EGF/FGF (blue) or BMP4/FGF (red). For all plots, the Y axis depicts the arcsinh transformed ratio versus column minimum for a particular antigen. A difference of 0.4 corresponds to an approximately 2-fold difference of total protein. Representative histograms of 24 (top) and 72 hours (bottom). For all plots, error bars contact the maximum and minimum values. For all antigens, at 24 hours N = 10 for EGF/FGF, N = 9 for BMP4/FGF; at 72 hours N = 12. (A) Quantification of levels of p-STAT3 S727 (24 hours: unpaired two-tailed t-test p = 0.0063; 72 hours: Mann-Whitney test p < 0.0001). (B) Quantification of levels of p-S6 S235/236 (24 hours: unpaired two-tailed t-test p = 0.8102; 72 hours: unpaired two-tailed t-test p = 0.3159). (C) Quantification of levels of p-S6 S240/244 (24 hours: unpaired two-tailed t-test p = 0.2393; 72 hours: unpaired two-tailed t-test p = 0.3461). (D) 20X representative images of E15.5 NSCs cultured for 6 days in media with EGF/FGF (left) or BMP4/FGF (right) stained for p-S6 S240/244 (white) and DAPI (blue). (E) Quantification of levels of p-4EBP1 T37/46 (24 hours: unpaired two-tailed t-test p = 0.0318; 72 hours: unpaired two-tailed t-test p < 0.0001). (F) 20X representative images of E15.5 NSCs cultured for 6 days in media with EGF/FGF (left) or BMP4/FGF (right) stained for p-4EBP1 T37/46 (white), Ki67 (yellow) and DAPI (blue). Scale bars for all 20X images = 50 µm.

### Decreases in mTOR-dependent phosphorylation of S6 are not sufficient to induce quiescence

Next, inhibition of these downstream components of mTOR signaling was tested for sufficiency in inducing quiescence. Rapamycin, a first generation mTOR inhibitor, has frequently been reported to inhibit the phosphorylation of S6 more effectively than that of 4EBP1/2 due to its specific binding location on mTOR (Choi et al., 1996; Fan et al., 2018). In cultures of embryonic day 15.5 NSCs, 30 nM of Rapamycin decreased levels of p-S6, but did not affect levels of p-4EBP1/2 after 4 hours (Fig 4A). After 24 hours in culture, p-S6 levels recovered and were not significantly different from vehicle treated levels (Extended Fig 4-1). Torkinib (PP242) is a second generation, dual mTORC1 and mTORC2 inhibitor that has been reported to inhibit phosphorylation of both S6 and 4EBP1/2 (Feldman et al., 2009), and was tested here in a dose response series ranging from 12.5 nM to 400 nM of Torkinib (Fig 4A). At all concentrations tested, 4-hour treatment with Torkinib decreased phosphorylation of S6 at both the mTOR-dependent S240/244 and alternate S235/236 residues. However, phosphorylation of 4EBP1/2 remained unaffected at all concentrations tested. Following acute (4 hour) and extended (24 hour) treatment with 400 nM Torkinib, levels of p-STAT3 S727 similarly did not decrease (Fig 4B). Additional second generation mTOR inhibitors and inhibitors of specific cap-dependent translation components were tested for their ability to decrease levels of p-4EBP1/2 in cultures; however, none had this ability (Extended Fig 4-1). Torkinib was therefore used as an S6-selective inhibitor to determine whether inhibition of phosphorylation of S6, but not 4EBP1/2, would be sufficient to induce quiescence. While levels of p-S6 S235/236 (Fig 4C) and p-S6 S240/244 (Fig 4D) initially decreased following 4 hour treatment with Torkinib, by 24 hours, both proteins had overcome the inhibition and were not different compared to vehicle treated cells. Phosphorylation of 4EBP1/2 was unaffected by treatment with Torkinib at both 4 and 24 hours (Fig 4E). Neither length of treatment affected levels of Ki67 (Fig 4F). While there was a small increase in the percentage of cells in G0/G1 with Torkinib treatment compared to vehicle after 24 hours of treatment (averages of 75.03% Torkinib vs 72.67% vehicle, N=9), there was not a significant decrease in the percent of cells in the S (7.73% vs 7.90%) or G2/M phases (16.01% vs. 15.06%) of the cycle with treatment (Fig 4G). Taken together, these data indicated inhibition of S6 phosphorylation alone was not sufficient to induce quiescence. Given that levels of p-4EBP1/2 decreased upon entry into quiescence, inhibition of p-4EBP1/2 was next tested for sufficiency to induce quiescence.

**Figure 4:**
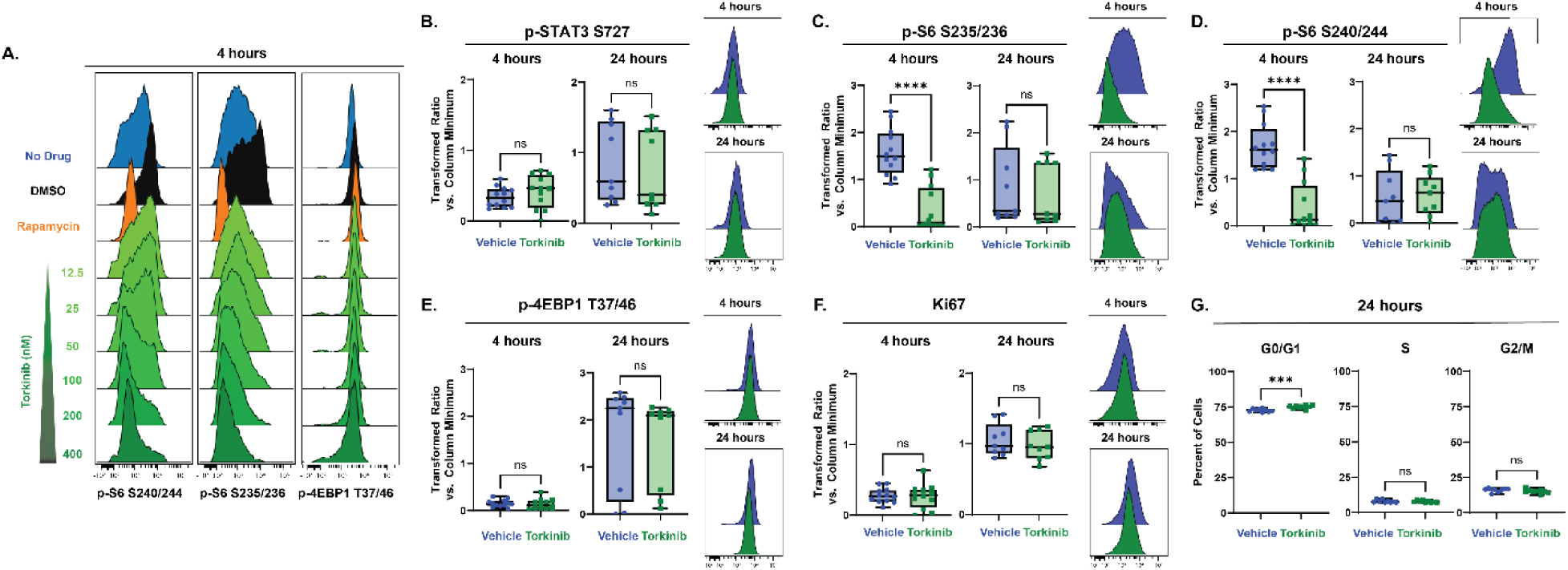
Decreases in mTOR-dependent phosphorylation of S6 are not sufficient to induce quiescence. (A) Representative histograms downstream effectors of mTOR from a dose-response experiment comparing E15.5 NSCs untreated (blue) to E15.5 NSCs treated with DMSO (0.06%, black), Rapamycin (30 nM, orange), or Torkinib (12.5 – 400 nM, green). Quantification of proliferation markers and effectors downstream of mTOR in E15.5 NSC cultures treated for 4 (left in plots) and 24 (right in plots) hours with media vehicle (1X PBS, blue) or 400 nM Torkinib (green). For all plots, the Y axis depicts the arcsinh transformed ratio versus column minimum for a particular antigen. A difference of 0.4 corresponds to an approximately 2-fold difference of total protein. Representative histograms of 4 (top) and 24 hours (bottom). For all plots, error bars contact the maximum and minimum values. For all antigens, at 4 hours, N = 12; at 24 hours, N =9. (B) Quantification of levels of p-STAT3 S727 (4 hours: unpaired two-tailed t-test p = 0.3123; 24 hours: unpaired two-tailed t-test p = 0.6878). (C) Quantification of levels of p-S6 S235/236 (4 hours: unpaired two-tailed t-test p < 0.0001; 24 hours: Mann-Whitney test p = 0.5457). (D) Quantification of levels of p-S6 S240/244 (4 hours: unpaired two-tailed t-test p < 0.0001; 24 hours: unpaired two-tailed t-test p = 0.8272). (E) Quantification of levels of p-4EBP1 T37/46 (4 hours: unpaired two-tailed t-test p = 0.5915; 24 hours: unpaired two-tailed t-test p = 0.8275). (F) Quantification of levels of Ki67 (4 hours: unpaired two-tailed t-test p = 0.8409; 24 hours: unpaired two-tailed t-test p = 0.4361). (G) Percent of E15.5 NSC cultures treated for 24 hours 400 nM Torkinib in each phase of the cell cycle. (N = 9; G0/G1 phase unpaired two-tailed t-test p = 0.0005, S phase unpaired two-tailed t-test p = 0.7134, G2/M unpaired two-tailed t-test p = 0.1695).

### Decreases in mTOR-dependent phosphorylation of 4EBP1/2 are sufficient to induce quiescence

RapaLink-1, a third generation mTOR inhibitor synthesized from Rapamycin and an ATP-competitive inhibitor of mTOR, MLN0128 (Rodrik-Outmezguine et al., 2016), has previously been shown to inhibit phosphorylation of 4EBP1/2 in mouse brain tissue (Zhang et al., 2022). Following 24 and 72 hours of treatment with 10 nM RapaLink, E15.5 NSCs were analyzed for readouts of the mTOR pathway. Levels of p-STAT3 S727 were unaffected compared to vehicle at 24 hours, but after 72 hours of treatment with RapaLink, the values were slightly decreased compared to vehicle (Fig 5A). Levels of both p-S6 S235/236 (Fig 5B) and p-S6 S240/244 (Fig 5C) decreased following 24 hours of treatment with RapaLink and stayed low compared to vehicle after 72 hours of treatment, contrasting with the transient effects of Torkinib. Levels of p-4EBP1/2 decreased compared to vehicle following 24 hours of treatment with RapaLink and remained decreased after 72 hours (Fig 5D). After 24 hours of treatment with RapaLink, levels of Ki67 were significantly decreased compared to vehicle and remained so at 72 hours (Fig 5E). After 24 hours of treatment, RapaLink treated cells had an increased percentage of cells in the G0/G1 phases of the cell cycle (averages of 78.63% with RapaLink vs. 67.42% with vehicle, N=7) and decreased percentages of cells in the S (12.27% vs. 18.71%) and G2/M phases (5.98% vs. 11.50%) of the cell cycle compared to vehicle treated cells. Taken together, these data indicate that inhibition of 4EBP1/2 is sufficient to induce quiescence in cultured embryonic NSCs.

**Figure 5:**
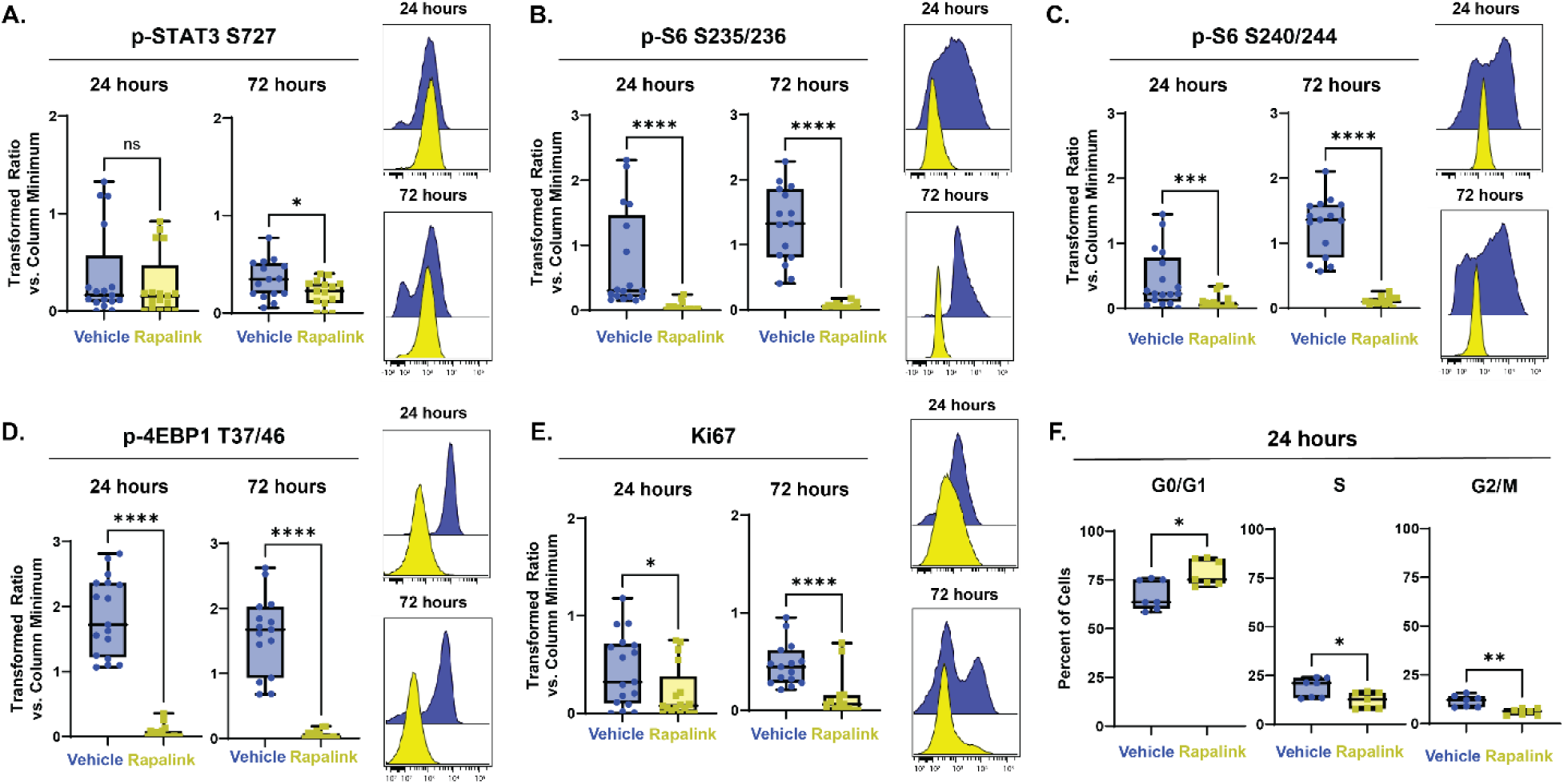
Decreases in mTOR-dependent phosphorylation of 4EBP1/2 are sufficient to induce quiescence. Quantification of proliferation markers and effectors downstream of mTOR in E15.5 NSC cultures treated for 24 (left in plots) and 72 (right in plots) hours with media vehicle (1X PBS, blue) or 10 nM RapaLink (yellow). For all plots, the Y axis depicts the arcsinh transformed ratio versus column minimum for a particular antigen. A difference of 0.4 corresponds to an approximately 2-fold difference of total protein. Representative histograms of 24 (top) and 72 hours (bottom). For all plots, error bars contact the maximum and minimum values. For all antigens, at 24 hours N = 17; at 72 hours N = 15. (A) Quantification of levels of p-STAT3 S727 (24 hours: unpaired two-tailed t-test p = 0.4550; 72 hours: unpaired two-tailed t-test p = 0.0330). (B) Quantification of levels of p-S6 S235/236 (24 hours: Mann-Whitney test p < 0.0001; 72 hours unpaired two-tailed t-test p < 0.0001). (C) Quantification of levels of p-S6 S240/244 (24 hours: Mann-Whitney test p = 0.0009; 72 hours: unpaired two-tailed t-test p < 0.0001). (D) Quantification of levels of p-4EBP1 T37/46 (24 hours: Mann-Whitney test p < 0.0001; 72 hours: unpaired two-tailed t-test p < 0.0001). (E) Quantification of levels of Ki67 (24 hours: unpaired two-tailed t-test p = 0.0433; 72 hours: Mann-Whitney test p < 0.0001). (F) Percent of E15.5 NSC cultures treated for 24 hours 10 nM RapaLink in each phase of the cell cycle. (N = 7, G0/G1 phase unpaired two-tailed t-test p = 0.0126, S phase unpaired two-tailed t-test p = 0.0223, G2/M unpaired two-tailed t-test p = 0.0011).

### Quiescence entry and mTOR response do not differ by dorsoventral position

Prior studies of postnatal cells found that NSCs from the ventral V-SVZ exhibited higher per-cell phosphorylation of mTORC1 targets than their dorsal counterparts (Rushing et al., 2019). To determine whether embryonic cells also had differential mTOR signaling based on cellular positioning, NSCs from the developing cortex (dorsal NSCs) and cells from the ganglionic eminences (ventral NSCs) were cultured separately and the various mTOR readouts were assessed before and after quiescence induction. In cultures maintained in media containing EGF/FGF-2, no uniform differences between dorsal and ventral NSCs in any of the readouts were observed across multiple studies. Following 72 hours of treatment with BMP4, Ki67 was decreased, and VCAM1 increased, in ventral NSCs grown in media containing BMP4/FGF-2 compared to controls (Fig 6A and 6B). These data suggest that exposure to BMP4 had a comparable effect inducing quiescence in both ventral NSCs and dorsal NSCs. Following 72 hours of BMP4 exposure, median levels of p-STAT3 S727 (Fig 6C) and p-4EBP1/2 T37/46 (Fig 6F) both decreased compared to ventral cells grown in EGF/FGF-2 media as they did in dorsal cells exposed to BMP4. However, as in dorsal NSCs, in ventral NSCs, levels of p-S6 S235/236 (Fig 6D) and p-S6 S240/244 (Fig 6E) did not change following 72 hours in culture with BMP4.

**Figure 6:**
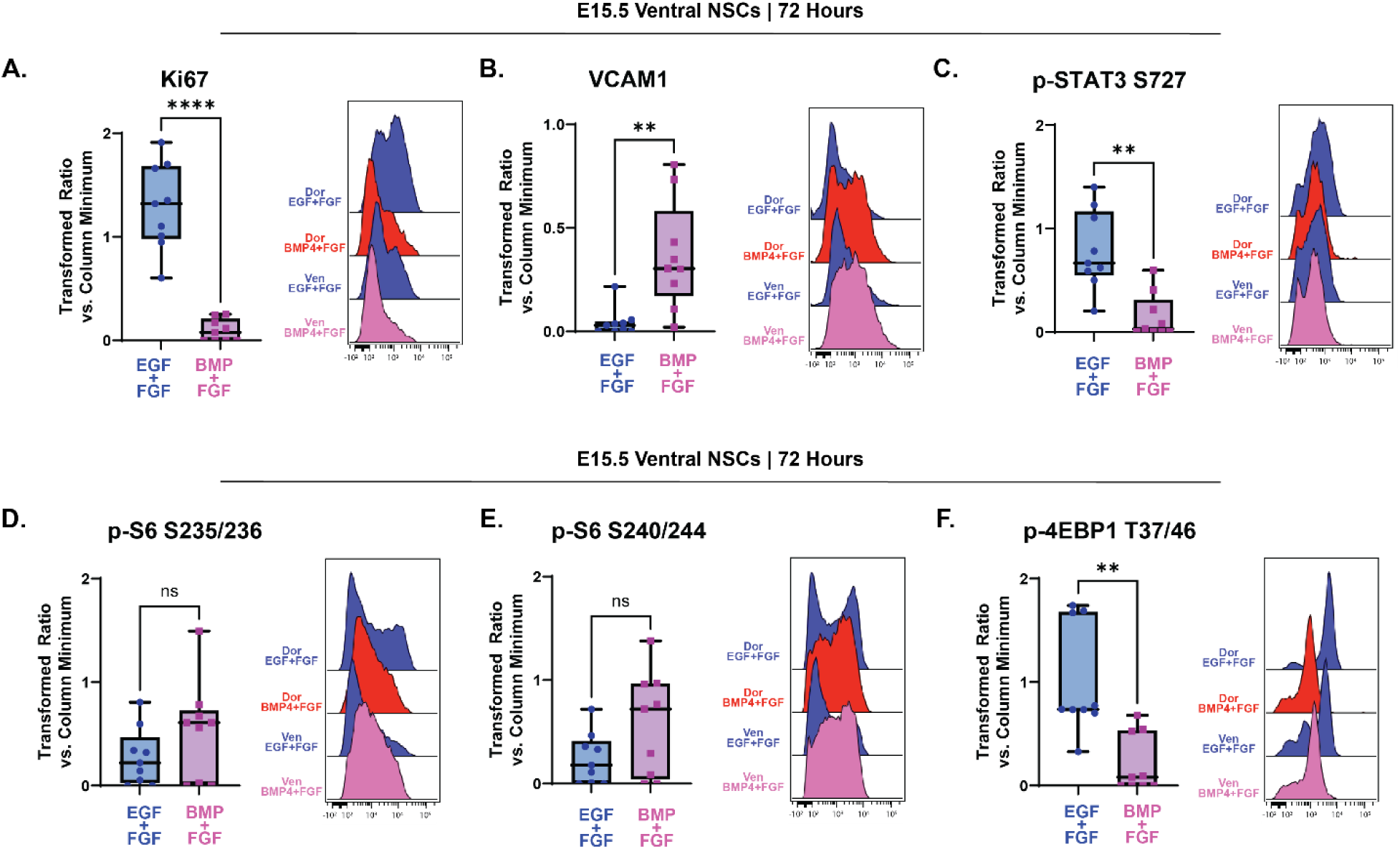
Quiescence entry and mTOR response do not differ by dorsoventral position. Quantification of effectors downstream of mTOR in E15.5 NSC ventral cultures grown 72 hours with media containing EGF/FGF (blue) or BMP4/FGF (pink). For all plots, the Y axis depicts the arcsinh transformed ratio versus column minimum for a particular antigen. A difference of 0.4 corresponds to an approximately 2-fold difference of total protein. Representative histograms comparing E15.5 dorsal and ventral NSC cultures grown in EGF/FGF media or BMP4/FGF media for 72 hours. For all plots, error bars contact the maximum and minimum values. For all antigens, N = 9. (A) Quantification of levels of Ki67 (paired two-tailed t-test p < 0.0001). (B) Quantification of levels of VCAM1 (Wilcoxon matched-pairs signed rank test p = 0.0039). (C) Quantification of levels of p-STAT3 S727 (paired two-tailed t-test p = 0.0015). (D) Quantification of levels of p-S6 S235/236 (paired two-tailed t-test p = 0.1707). (E) Quantification of levels of p-S6 S240/244 (paired two-tailed t-test p = 0.1871). (F) Quantification of levels of p-4EBP1 T37/46 (paired two-tailed t-test p = 0.0053).

## Discussion

In both the ventricular-subventricular zone and the dentate gyrus of the hippocampus, temporary quiescence of a subset of embryonic neural stem cells determines the capacity of the adult neurogenic niche through preservation of self-renewing stem cells that later activate in the adult. Lineage tracing work in rodents has shown extensive consequences of altering prenatal quiescence entry in both stem cell niches, including premature depletion of the postnatal stem cell population and developmental and learning defects (D. Y. Wang et al., 2020; Hu et al., 2017; Kokovay et al., 2012). This quiescence entry is thus essential to produce adult neural stem cells and support adult neurogenesis in both the V-SVZ and dentate gyrus. However, the mechanisms regulating the initiation of this essential process remain poorly understood (Urbán, 2022; Urbán et al., 2019).

mTOR has been shown to regulate the balance of activation and quiescence in other stem cell populations (Cho & Hwang, 2012; Nieto-González et al., 2019; Rodgers et al., 2014). In postnatal NSCs, mTOR also regulates preferential translation of specific mRNA transcripts as stem cells are activated to divide — that is, upon exit from quiescence (Baser et al., 2019; Rossi et al., 2021). The data shown here illuminate a role for this kinase in embryonic NSC quiescence. While the sufficiency of changes in p­4EBP1/2 for initiation of quiescence entry is demonstrated here, the necessity and sufficiency of increased p-4EBP1/2 for quiescence exit in these cells remain to be explored. Nutrient sensing and PI3K/Akt signaling upstream of mTOR have been proposed as a regulator of NSC quiescence exit (Chell & Brand, 2010). Whether mTOR-mediated phosphorylation of 4EBP1/2 is a regulator of both quiescence entry and exit or how it may work coordinately with other signaling pathways well known to be involved in quiescence, such as Notch signaling, is an area of ongoing study.

These data demonstrate that two primary downstream effectors of mTORC1, p-S6 S240/244 and p-4EBP1/2 T37/46, have distinct patterns of expression throughout neurogenesis and rarely appear in the same cell (fewer than 1% of cells at E15 and E17). Cells positive for p-4EBP1/2 line the most apical portion of the developing telencephalon, the ventricular zone, and their abundance decreases with age, while cells positive for p-S6 predominate in the subventricular zone, intermediate zone, and cortical plate and their abundance increases with age. The finding that p-4EBP1/2 and p-S6 were so rarely expressed in the same cell indicates that though the mTOR pathway is active in both cells, through a yet unidentified regulatory mechanism, only one signaling effector is being phosphorylated. While it has been reported that different ligands (such as amino acids, insulin, and growth factors) activating different upstream receptors (including the epidermal growth factor receptor, the fibroblast growth factor receptor, and the insulin receptor) can result in various downstream signaling responses (Sparta et al., 2021), it remains to be explored whether this type of mechanism is responsible for the differing patterns of phosphorylation of these two key mTORC1 effectors in NSCs.

The use of BMP4 in embryonic NSC cultures resulted in decreased expression of Ki67, increased expression of VCAM1, an increased percentage of cells in the G0/G1 phase of the cell cycle and decreased percentages of cells in the S and G2/M phases of the cell cycle. Cells exposed to BMP4 did not enter senescence (data not shown). Upon quiescence entry, levels of p-4EBP1/2, but not p-S6, decreased in cultures derived from the developing dorsal region. The wider variance seen in S6 signaling may reflect its role in additional biological processes, such as regulating cell size (Hartman et al., 2013; Magnuson et al., 2012; Montagne et al., 1999; Ruvinsky & Meyuhas, 2006). In concordance with the effects of BMP4, inhibition of p-4EBP1/2, but not p-S6, was sufficient to induce quiescence entry. Importantly, while levels of another mTOR downstream effector, p-STAT3, decreased in both dorsal and ventral NSCs exposed to BMP4, it did not decrease upon entry into quiescence after 24 hours with RapaLink treatment while cells had already begun to exit the cell cycle and quiesce. The opposing relationships of BMP and STAT3 signaling through mTOR to influence stem cell fate have been previously described (Rajan et al., 2003). The data here indicate the inhibition of phosphorylation of STAT3 is not necessary for quiescence entry and may be a secondary consequence, whereas mTOR-dependent phosphorylation of 4EBP1/2 is a regulator of embryonic NSC quiescence entry.

While p-4EBP1/2 positive cells were nearly always dividing and co-expressed Ki67, p-S6 positive cells co-expressed Ki67 only 30% of the time, on average. This pattern may offer insight into the independent biological functions that activation of each signaling protein triggers and the consequences on stem cell proliferation, self-renewal, and differentiation those functions have. It has been widely reported that as stem cells differentiate and migrate tangentially away from the ventricular surface, translation is suppressed. This suppression of translation, and regulation of the process by mTOR, has been hypothesized to be a mechanism of regulating stem cell fate (reviewed in R. Wang & Amoyel, 2022). mTOR-mediated translation of specific transcripts — or lack thereof — during key periods of neurogenesis has been shown to regulate cellular differentiation and neuronal subtype specification (Harnett et al., 2022; reviewed in Statoulla et al., 2021). The data presented here support this hypothesis, as p-4EBP1/2, a key regulator of translation, decreases with increasing distance from the ventricular surface and as embryonic development proceeds. These data also offer a new insight into the role of p-S6 in stem cell differentiation to be explored in future studies, as p-S6 is markedly absent at the ventricular surface but its abundance increases with increasing distance from the ventricular surface.

An important implication of these data is that each downstream effector of mTOR should be investigated independent of the other signaling molecules in each cell type. mTOR signaling has been implicated as a proposed regulatory mechanism in multiple aspects of neural development and a variety of diseases of the nervous system (Andrews et al., 2020; Avet-Rochex et al., 2014; Costa-Mattioli & Monteggia, 2013; D’Gama et al., 2017; Hartman et al., 2013; Ka et al., 2014; Lee, 2015; Licausi & Hartman, 2018; D. Liu et al., 2018; Mahoney et al., 2016; Maierbrugger et al., 2020; Musah et al., 2020; Paliouras et al., 2012; Rushing et al., 2019; Tee et al., 2016; Tyler et al., 2009; Wahl et al., 2014; Zeng et al., 2009). Often, however, only a single residue on p-S6 — either S235/236 or S240/244 — is reported as a representative readout of total mTOR kinase activity. The data here demonstrate that the multiple signaling effectors of mTORC1 behave independently in tissue, *in vitro,* and in response to different pharmacological modulators. More broadly, as multiple generations of mTOR inhibitors enter clinical trials, the use of agents that more effectively inhibit phosphorylation of both S6 and 4EBP1/2 are likely to have broader effects on normal neural development, and cortical hyperplasias, than their predecessors.

Three different generations of mTOR inhibitors were tested here for their ability to inhibit phosphorylation of 4EBP1/2. While the inhibitors tested have all been reported to decrease p-4EBP1/2 levels in cell lines and *in vitro* assays, only the third-generation bivalent inhibitor, RapaLink-1, was able to decrease levels of p-4EBP1/2 in embryonic neural stem cell cultures. Rapamycin, a first-generation inhibitor, multiple second-generation “Tork” inhibitors, and a eukaryotic initiation factor inhibitor all failed to decrease levels of p-4EBP1/2. The data presented here may indicate cell type-specific mechanisms regulating mTOR signaling and susceptibility to inhibition. This finding has potential implications for clinical use, where mTOR inhibitors are often prescribed for a variety of diseases. First generation mTOR inhibitors (rapalogs) are often prescribed for pediatric patients with “mTORopathies,” a debilitating class of neurodevelopmental disorders. Patients with tuberous sclerosis complex, one such mTORopathy wherein patients have tumors throughout the entire body, are regularly prescribed the rapalog everolimus to control seizures and limit brain tumor growth (Cavalheiro et al., 2021; Feliciano, 2020; Franz, 2011; Karalis & Bateup, 2021; Overwater et al., 2019). This work may indicate that only the S6 “arm” of the mTOR pathway is inhibited by rapalog treatment and may offer insight as to why such treatments are not cytotoxic, but merely cytostatic. These data support the testing of improved mTOR inhibitors that more effectively inhibit phosphorylation of 4EBP1/2 in neural stem cells, but also raise concern that this targeting may incur additional side effects.

An additional area of future study is the comparison to the human brain. The day 10 neurospheres derived from human iPSCs presented here suggest the independent phosphorylations of S6 and 4EBP1/2 also occur in human cells. Outer radial glial cells, a cell type unique to the human brain hypothesized to be a cell of origin in disease, have been reported to have increased mTOR activity, as measured by p-S6, compared to other types of cells (Andrews et al., 2020; Nowakowski et al., 2017). Embryonic neural stem cells have been hypothesized to be the cell of origin for some of the brain tumor types found in tuberous sclerosis complex (Blair et al., 2018; Eichmüller et al., 2022; Hang et al., 2017; Hewer & Vajtai, 2015; Rushing et al., 2019). However, more work is needed to determine whether there are changes in mTOR-dependent phosphorylation of 4EBP1/2 upon quiescence entry in human NSCs. Future studies may investigate how consequences of altered quiescence entry, particularly in the context of disease, may affect a stem cell’s lineage and fate.

## Supporting information

Extended Figures_Compiled

## Acknowledgements

The authors would like to acknowledge the members of the Ihrie and Irish laboratories at Vanderbilt University and the members of the Ess laboratory at Vanderbilt University Medical Center for their helpful feedback on experiments and data interpretation throughout the duration of this project. Additionally, we acknowledge the staff of the Translational Pathology Shared Resource (supported by the NIH P30 CA68485 grant), Cell Imaging Shared Resource (supported by NIH grants CA68485, DK20593, DK58404, DK59637 and EY08126), Flow Cytometry Shared Resource (supported by the NIH P30 CA68485 grant and the Vanderbilt Digestive Disease Research Center grant DK058404), and Jose Maldonado at the Neurovisualization Lab at Vanderbilt University and Vanderbilt University Medical Center for their work and expertise in completing these experiments. We gratefully acknowledge the work of Moesha Parsons in sectioning mouse brains and Michael Martland for his work testing antibodies for use in immunostaining.

---

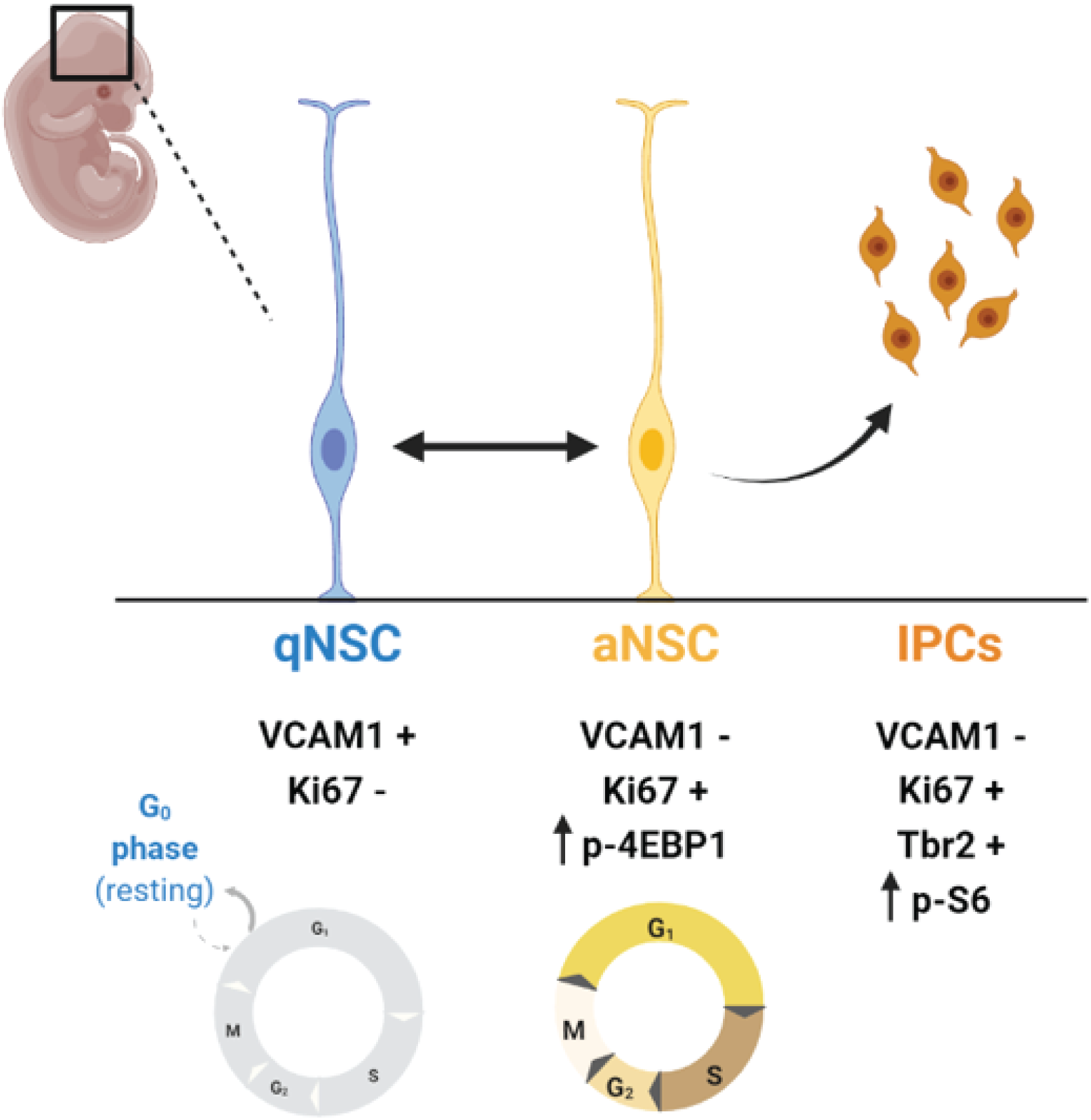

## Notes

Funding Information This research was supported by the following funding resources: NIH T32 GM07628 (LCG), NIH F31 NS120608 (LCG), NIH T32 HD007502 (MBLC), NIH F31 HD106890 (SRS), NIH R01 DK106476 (RBS), NIH R01 NS118580 (KCE, RAI, MBLC), Ben & Catherine Ivy Foundation (RAI), NIH R01 NS096238 (R.A.I., J.M.I.), NIH R01 CA226833 (J.M.I.), NIH U54 CA217450 (J.M.I.), the Michael David Greene Brain Cancer Fund (R.A.I., J.M.I.), the Southeastern Brain Tumor Foundation (R.A.I., J.M.I.), and the Vanderbilt-Ingram Cancer Center (VICC, P30 CA68485).

https://zenodo.org/record/7641606

### Competing Interest Statement

The authors have declared no competing interest.

