## Extended Figures_Compiled for "Dephosphorylation of 4EBP1/2 Induces Prenatal Neural Stem Cell Quiescence"

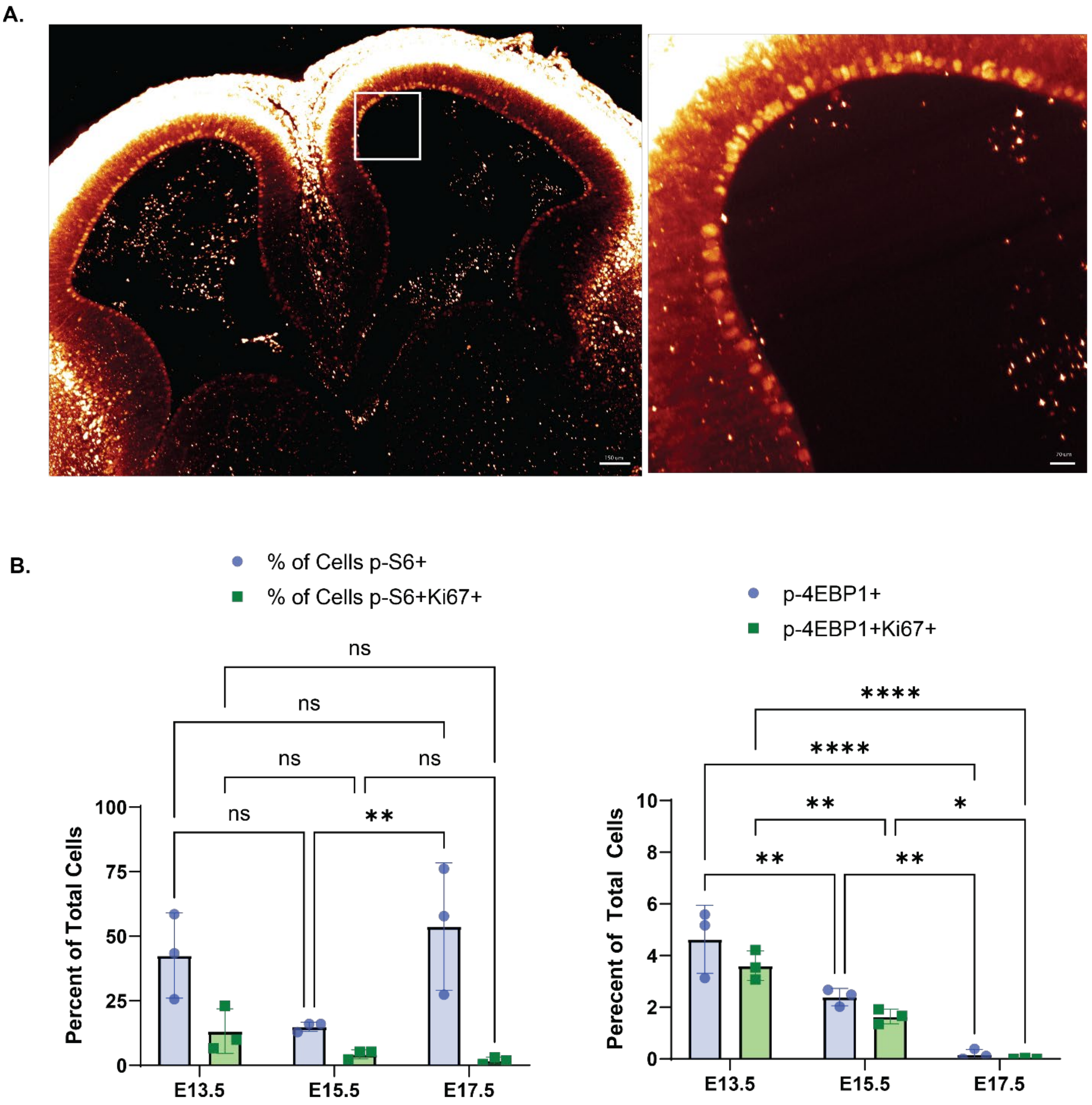


**Extended Figure 1-1: Additional staining of mTOR during neurogenesis.** (A) iDISCO+-tissue cleared whole brain from an E13.5 mouse embryo (15X) Ki67 positive cells lining the surface of the ventricular-subventricular zone tissue and in the developing cortex (inset). (B) Plots showing the quantification of mTOR signaling throughout neurogenesis. 2 way ANOVA with Tukey's multiple comparisons test: percent of p-S6 positive cells: E13.5 versus E15.5 $p = 0.0501$, E15.5 versus E17.5 $p = 0.0071$, E13.5 versus E17.5 $p = 0.5377$; percent of p-S6 and Ki67 positive cells: E13.5 versus E15.5 $p = 0.6704$, E15.5 versus E17.5 p =

0.9697, E13.5 versus E17.5 $p = 0.5301$). (C) The percent of all cells positive for p-4EBP1 T37/46 (green) and cells co-positive for p-4EBP1 and Ki67 (red) (bottom). (2 way ANOVA with Tukey's multiple comparisons test: percent of p-4EBP1 positive cells: E13.5 versus E15.5 $p = 0.0022$, E15.5 versus E17.5 $p = 0.0023$, E13.5 versus E17.5 $p < 0.0001$; percent of p-4EBP1 and Ki67 positive cells: E13.5 versus E15.5 $p = 0.0056$, E15.5 versus E17.5 $p = 0.0184$, E13.5 versus E17.5 $p < 0.0001$). Error bars represent standard deviation.

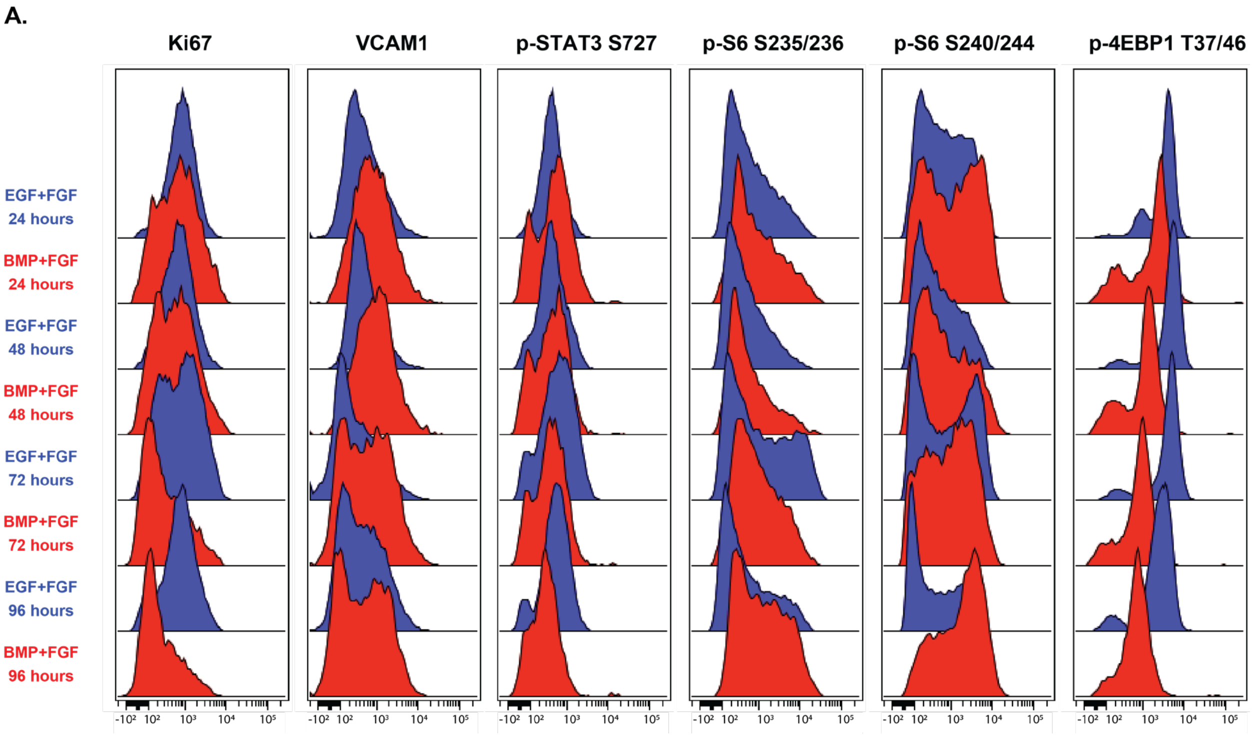


**Extended Figure 2-1: Quiescence entry time course.** (A) Representative histograms of proliferation markers and effectors downstream of mTOR in E15.5 NSCs cultured for 24, 48, 72, and 96 hours with media containing either EGF/FGF (blue) or BMP4/FGF (red).

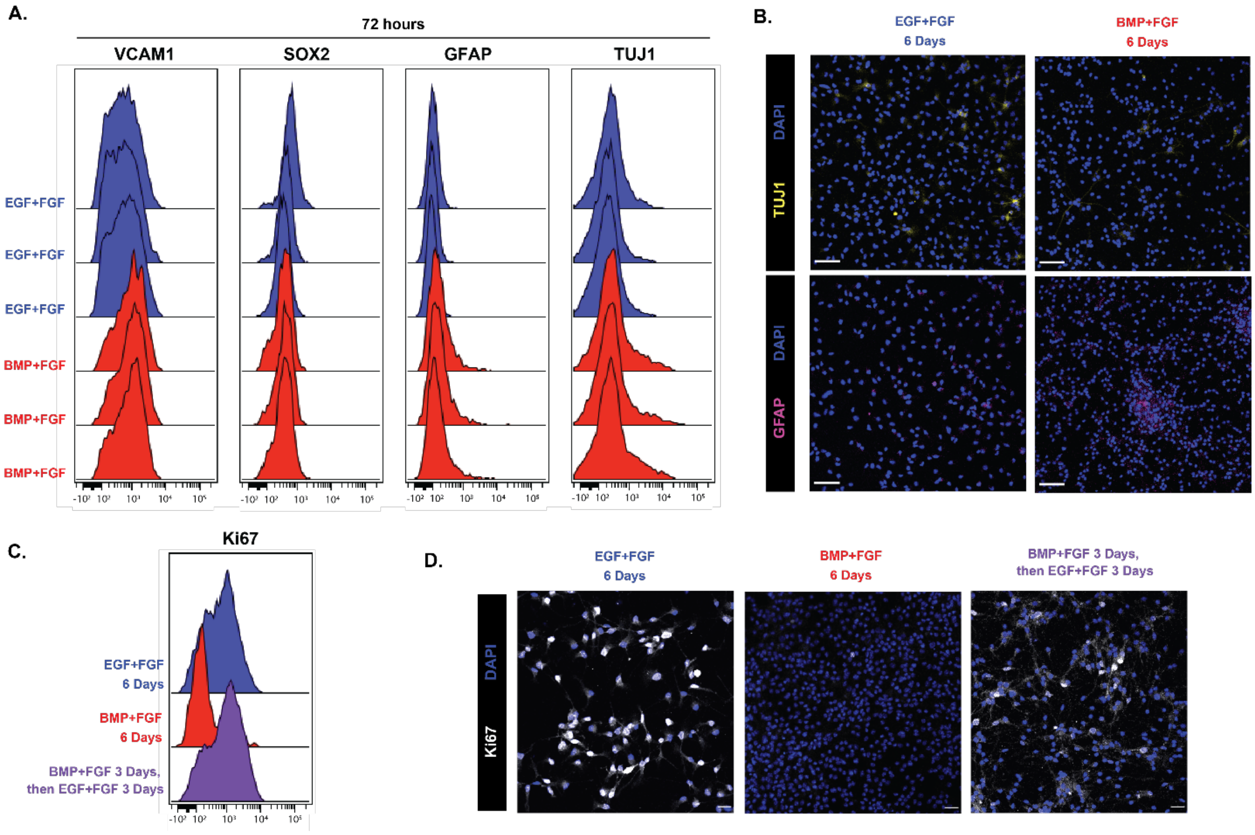


**Extended Figure 2-2: BMP4 transiently induces quiescence, but not differentiation.** (A) 3 replicates of E15.5 NSCs cultured for 72 hours with media containing either EGF/FGF (blue) or BMP4/FGF (red) stained for VCAM1, Sox2, GFAP, and TUJ1. (B) 20X representative images of E15.5 NSCs cultured for 6 days with media containing either EGF/FGF (left) or BMP4/FGF (right) stained for TUJ1 (top, yellow), GFAP (bottom, pink), and DAPI (blue). Scale bars = 50 µm. (C) Histogram overlays of E15.5 NSCs cultured for 6 days with media containing EGF/FGF (blue), 6 days with media containing BMP4/FGF (red), and 3 days in BMP4/FGF followed by 3 days back in EGF/FGF (purple). (D) Representative 20X images of E15.5 NSCs cultured for 6 days with media containing EGF/FGF (left), 6 days with media containing BMP4/FGF (center), and 3 days in BMP4/FGF followed by 3 days back in EGF/FGF (right) stained for Ki67 (white) and DAPI (blue). Scale bars = 20 µm.

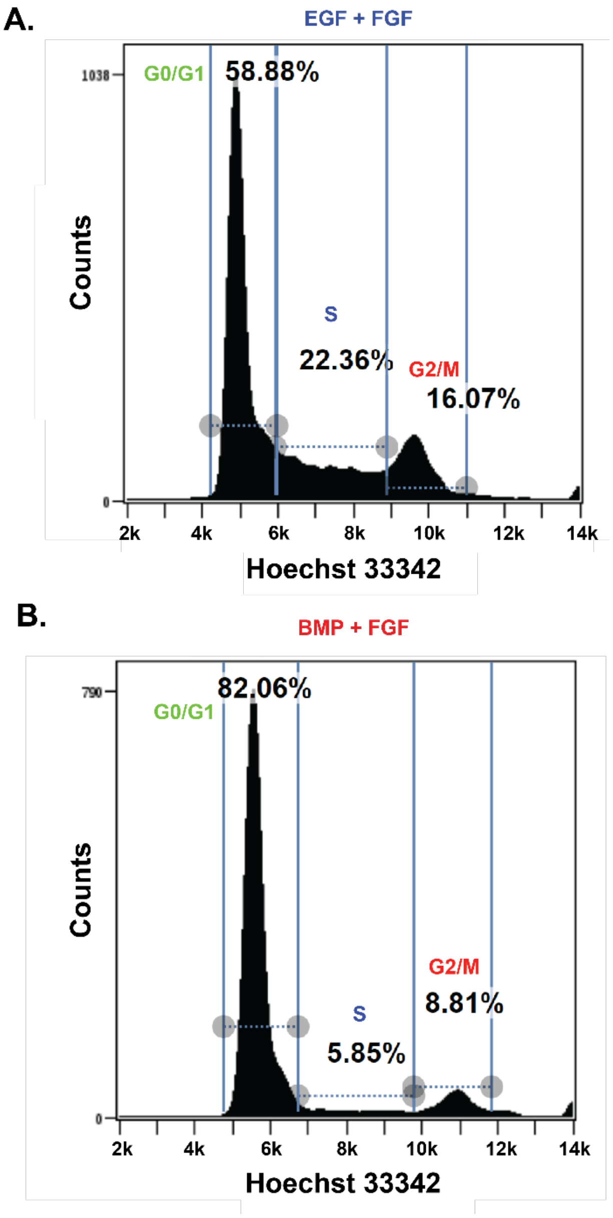


**Extended Figure 2-3: Gates for determining phases of cell cycle.** (A) Histogram of E15.5 NSCs cultured for 24 hours in media containing EGF/FGF stained with Hoechst 33342. The G0/G1 gate depicts 2n cells and the G2/M gate depicts 4n cells. (B) Histogram of E15.5 NSCs cultured for 24 hours in media containing BMP4/FGF stained with Hoechst 33342. The G0/G1 gate depicts 2n cells and the G2/M gate depicts 4n cells, with cells in S phase in between these two populations.

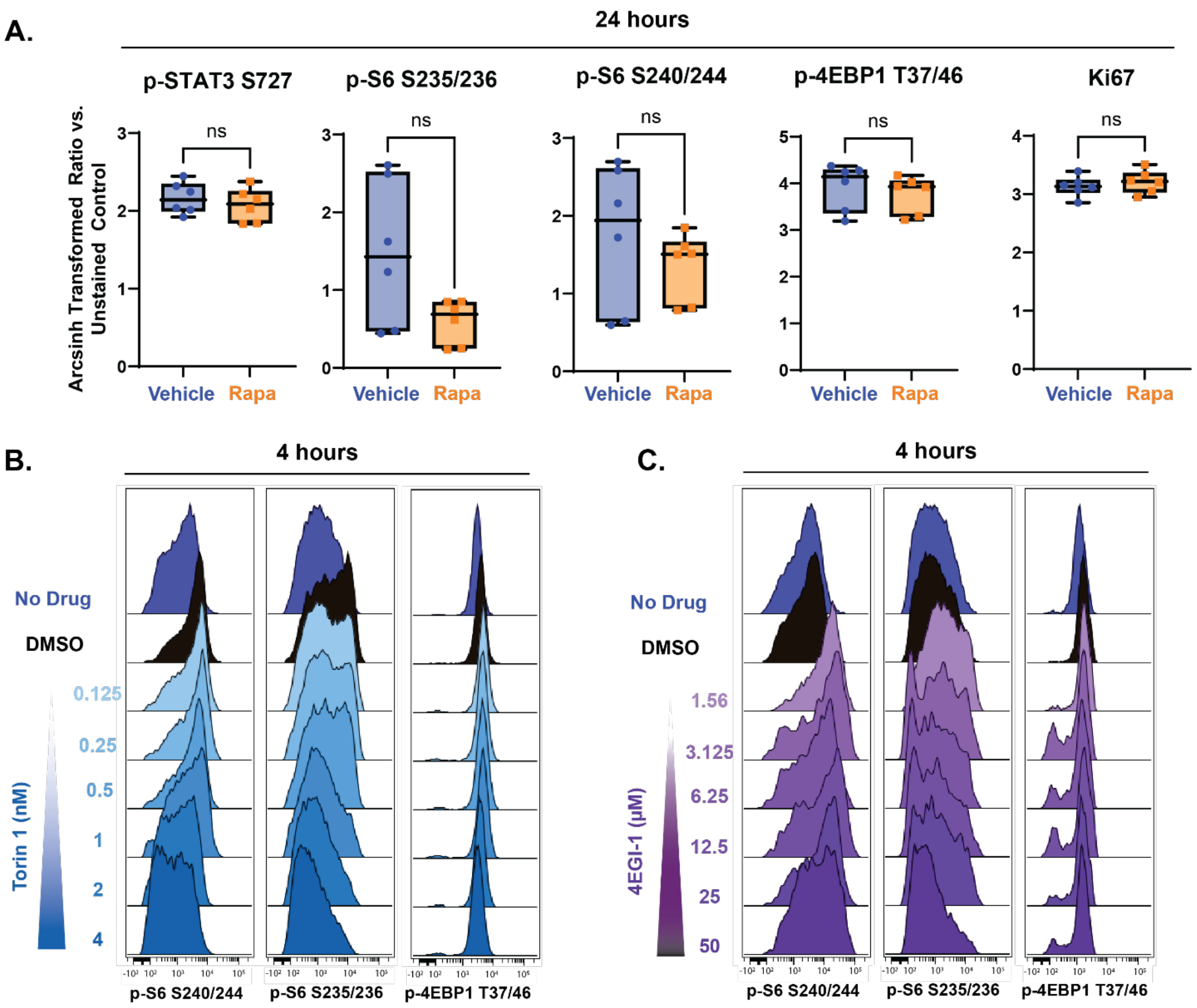


**Extended Figure 4-1: Dose response tests of various mTOR inhibitors.** (A) Quantification of proliferation markers and effectors downstream of mTOR in E15.5 NSC cultures treated for 24 hours with media vehicle (1X PBS, blue) or 30 nM Rapamycin (orange). For all plots, the Y axis depicts the arcsinh transformed ratio versus column minimum for a particular antigen. A difference of 0.4 corresponds to an approximately 2-fold difference of total protein. For all plots, error bars contact the maximum and minimum values. N = 9. p-STAT3 S727 (unpaired two-tailed t-test $p = 0.7921$), p-S6 S235/236 (unpaired two-tailed t-test $p = 0.0335$), p-S6 S240/244 (unpaired two-tailed t-test $p = 0.3306$), p-4EBP1 (unpaired two-tailed t-test $p = 0.9630$), and Ki67 (unpaired two-tailed t-test $p = 0.6177$). (B) Representative histograms downstream effectors of mTOR from a dose-response experiment comparing E15.5 NSCs untreated (blue) to E15.5

NSCs treated with DMSO (0.06%, black) or Torin 1 (0.125 – 4 nM, blue). (C) Representative histograms downstream effectors of mTOR from a dose-response experiment comparing E15.5 NSCs untreated (blue) to E15.5 NSCs treated with DMSO (0.1%, black) or 4EGI-1 (1.56 – 50 nM, blue).
